# Disorder in Ca^2+^ Release Unit Locations Confers Robustness but Cuts Flexibility of Heart Pacemaking

**DOI:** 10.1101/2021.11.19.469309

**Authors:** Anna V. Maltsev, Victor A. Maltsev

**Author notes:** **Corresponding author:** Anna V. Maltsev.

## Abstract

Excitation-contraction coupling kinetics are dictated by the rate and rhythm of the excitations generated by sinoatrial-nodal cells. These cells generate local Ca releases (LCRs) that activate Na/Ca exchanger current, which accelerates diastolic depolarization and determines the rate and rhythm of the excitations. The LCRs are generated by clusters of ryanodine receptors, Ca release units (CRUs), residing in the sarcoplasmic reticulum. While the spatial CRU distribution in pacemaker cells exhibits substantial heterogeneity, it remains unknown if it has any functional importance. Using numerical modeling, here we showed that with a square lattice distribution of CRUs, Ca-induced-Ca-release propagation during diastolic depolarization is insufficient for pacemaking within a broad lower range of realistic *I*_*CaL*_ densities. Allowing each CRU to deviate from its original lattice position fundamentally changes the model behavior: during diastolic depolarization sparks propagate, forming LCRs observed experimentally. As disorder in the CRU positions increases, the CRU distribution exhibits larger empty spaces but simultaneously CRU clusters, as in Poisson clumping. Propagating within the clusters, Ca release becomes synchronized, increasing AP firing rate and reviving pacemaker function within lower *I*_*CaL*_ densities at which cells with lattice CRU distribution were dormant/non-firing. However, cells with fully disordered CRU positions cannot reach low firing rates and their β-adrenergic receptor stimulation effect was substantially decreased. Thus, order/disorder in CRU locations regulates Ca release propagation and could be harnessed by pacemaker cells to regulate their function. Excessive disorder is expected to limit heart rate range that may contribute to heart rate range decline with age and in disease.

**Summary:** The present numerical modeling study shows that disorder in locations of Ca release units in cardiac pacemaker cells has substantial functional impact by creating release clusters, similar to Poisson clumping, and opportunity of Ca release to propagate within the clusters.

## Introduction

Each excitation-contraction coupling cycle in the heart begins with the generation of rhythmic excitation in the sinoatrial node (SAN). To satisfy a given blood supply demand, cardiac muscle performance, defined by its state of contractile apparatus, Ca cycling proteins, and cell excitability must be in balance with the rate and rhythm of the excitation impulses generated by the SAN. Thus, the cardiac pacemaker function is a vital part of the excitation-contraction coupling that “sets the stage” for timely interactions for all further downstream mechanisms.

In turn, the generation of normal rhythmic cardiac impulses is executed via timely interactions within and among SAN pacemaker cells that involve a coupled signaling of both cell membrane ion channels and Ca cycling, dubbed a coupled-clock system (Maltsev and Lakatta, 2009; Lakatta et al., 2010). A key element of the system is sarcoplasmic reticulum (SR) that rhythmically generates diastolic local Ca releases (LCRs) via Ca release channels, ryanodine receptors (RyRs). The LCRs contribute to SAN cell pacemaker function via activation of Na/Ca exchanger (NCX) current (*I*_*NCX*_) that accelerates the diastolic depolarization (Huser et al., 2000; Bogdanov et al., 2001; Lakatta et al., 2010). Initially the role of Ca release in cardiac pacemaker function has been numerically studied in so-called “common pool” models (Kurata et al., 2002; Himeno et al., 2008; Maltsev and Lakatta, 2009; Imtiaz et al., 2010; Severi et al., 2012), in which local Ca dynamics was neglected and cell Ca release was approximated by a single variable. While such traditional simplified approach has yielded substantial progress in our understanding of Ca release role in pacemaker function (including modern theories of coupled-clock function (Maltsev and Lakatta, 2009) and ignition (Lyashkov et al., 2018)), it has fundamental limitations. Thus further progress requires a new type of modeling that would take into account local interactions (Maltsev et al., 2014). A general issue is that we still do not have a clear quantitative understanding of how the Ca clock and the coupled clock system emerge from the scale of molecules (such as RyRs) towards the whole cell function (Weiss and Qu, 2020). Thus how local and whole-cell signaling events that coordinate cell function emerge from stochastic openings of individual release channels remains an unresolved fundamental problem despite extensive investigation.

A concrete issue with common pool models is that they operate with Ca signals in sub- μM range, i.e. with at least two orders of magnitude lower concentrations than those Ca sparks and diastolic LCRs can actually reach (tens and even hundreds of μM in their amplitudes (Stern et al., 2013; Stern et al., 2014)). The molecules involved in pacemaker function (ion channels, exchangers, pumps, and Ca-sensitive enzymes), however, operate via the high local Ca concentrations rather than a whole-cell average. Common pool models thus are analogous to a mean-field theory approach in physics, which usually does recover faithfully qualitative information about system behavior, but fails to zone in on the critical parameter values.

While the membrane clock operates as a limit cycle oscillator (Kurata et al., 2012), the Ca-clock seems to operate by completely different mechanisms based on phase transitions (Maltsev et al., 2011) or criticality (Nivala et al., 2012; Weiss and Qu, 2020). RyRs are organized and operate in clusters of 10-150 channels (Greiser et al., 2020), known as Ca release units (CRUs) in all cardiac cells, including SAN cells (Maltsev et al., 2011; Stern et al., 2014). Ca release in ventricular myocytes occurs mainly via Ca sparks (defined as a single CRU Ca release) tightly controlled by L-type Ca channel openings (local control theory (Stern et al., 1997)), whereas LCRs in SAN cells consist of multiple sparks that emerge spontaneously via self-organization by means of positive feedback provided by Ca-induced Ca release (CICR). The synchronized CRU activation leads to oscillatory, phase-like transitions in SAN cells (Maltsev et al., 2011) that generate a net diastolic LCR signal that ultimately interacts with NCX and L-type Ca channels (Maltsev et al., 2013; Lyashkov et al., 2018) to ignite pacemaker action potentials (APs), which comprises the ignition theory of APs (Lyashkov et al., 2018). The probability that a Ca spark can “jump” in this way to activate its neighbor and form a propagating multi-spark LCR instead of remaining an isolated event depend on various parameters, including in particular the amplitude of the spark (sometimes called *I*_*spark*_ (Zhou et al., 2009; Maltsev et al., 2011)) and its nearest neighbor distance (Stern et al., 2014). The difficult and fascinating phenomenon that occurs in this system is that the area affected by propagation of an LCR is discontinuous in the probability of the Ca spark “jump.” This discontinuity is a phase-like transition first demonstrated in (Maltsev et al., 2011), and the exact parameter values at which it occurs is known as criticality. Thus, to understand the intrinsic mechanisms of SAN cell operation, it is important to quantitatively predict critical parameter values at which the system changes its operational paradigm from sparks to LCRs.

Initially, the LCRs were numerically studied with CRUs located in a perfect rectangular lattice (Maltsev et al., 2011; Maltsev et al., 2013). These studies showed that the ability of LCR to propagate regulates the size and the impact of the LCRs on the diastolic depolarization. However, RyR immunofluorescence exhibited notable disorder in the CRU distribution (Stern et al., 2014; Maltsev et al., 2016) and a more recent numerical model (Stern et al., 2014) was developed with two fixed sizes of CRUs. The larger CRUs were located in a rectangular lattice that lacked release propagation until smaller clusters were introduced in the CRU network providing bridges for release propagation among the larger CRUs. While this “invented” CRU is closer to realistic distribution and provided functionality to the SAN cell model, the total number of CRUs was not controlled and whether or not disorder in the CRU distribution per se is capable of impacting SAN cell function has not been examined and represents the specific aim of the present study.

Here we approach the problem via numerical model simulations of SAN cell function with the same number of identical CRUs, but different spatial CRU distributions. Specifically, we test how various degree of disorder (or noise) in CRU positions would influence SAN cell operation. Surprisingly, the least robust SAN cell function is achieved when CRU positions are distributed in the perfect rectangular lattice. As disorder in the CRU positions increases, nearest neighbor distances decrease, creating “shortcuts” for Ca release propagation resulting in AP firing rate increase and robust pacemaker function. The most robust function was achieved when CRUs were distributed uniformly randomly, i.e. independent of the perfect grid. However, the robust function comes at the cost of the chronotropic reserve: disorder-facilitated broad propagation of Ca release in the basal state substantially diminishes the unutilized propagation capability, which normally is available to accelerate AP firing rate during β-adrenergic receptor (βAR) stimulation. Thus, disorder in spatial CRU distribution provides a novel subcellular mechanism of cardiac pacemaker regulation that a SAN cell may utilize to achieve a perfect balance between robustness and flexibility.

## Methods

Our study of CRU interactions fully relies on the quality and predictive power of our numerical model of SAN cell. Therefore, as noted in Introduction, this aspect of the study merits particular attention. Common pool models, such as of Kurata et al. (Kurata et al., 2002) or Maltsev-Lakatta (Maltsev and Lakatta, 2009), is on the scale of a cell, lumping the contributions of all CRUs into one variable describing Ca release and one pool of junctional SR with fixed total SR volume. On the other hand, a 3D model of SAN cell developed by Stern et al. (Stern et al., 2014) featuring interactions of individual RyRs (i.e. RyR-based model, or Stern model for short) is on the scale of a CRU, as it provides too many fine details of intra-CRU Ca dynamics at a much higher computational cost and which are not so important for understanding Ca dynamics at the level of CRU to CRU.

Thus, an adequate model for our study would deal with Ca release at the level of CRU (as detected by confocal microscopy). We have previously developed such a model (CRU-based model, for short) featuring a two-dimensional array of stochastic, diffusively coupled CRUs located under cell membrane (Maltsev et al., 2011), and then added the full electrophysiological membrane clock to this model (Maltsev et al., 2013). While approximation of Ca release at the level of individual CRUs suits our study, here we performed a further major model update that allows for the model to predict the effect of CRU distribution on the AP firing rate in basal state and during β-AR stimulation in mechanistical setting of the “coupled clock” with a minimum number of independent variables (i.e. less uncertainties). The model structure is schematically illustrated in Fig. 1. The full details of the updated model are given in the Appendix. In short, we added the following critical features to our model:

1. We want to study interaction (synchronization) of CRU firing via CICR. However, a major weakness of our old model was that while it predicted the stochastic CRU activation beyond the refractory period, it could not predict the refractory period itself. Thus, it was not a mechanistic “coupled-clock” model because the refractory period of CRU firing Ca release (being the essence of the clock) was in fact an independent model parameter obtained as a direct read-off value from experimental data (Maltsev et al., 2011). After the refractory period, CRU was allowed to open with a given probability per unit time. The stochastic CRU firing generated a spark of a fixed duration that was another independent model variable tuned to experimental data. Here we modified the model to predict both the timing of spark activation and termination based on the present knowledge in this research area. A typical spark generated by our new model is shown in Fig. A1. Sparks are activated to release Ca when refilling junctional SR with Ca (via SERCA pumping and intra-SR diffusion) in diastole reaches a certain critical level as suggested in experimental studies (Vinogradova et al., 2010), numerical model simulations via common pool models (Maltsev and Lakatta, 2009; Imtiaz et al., 2010), RyR-based models (Stern et al., 2014), and in a recent theoretical study based on Ising formalism (Veron et al., 2021).
2. In the previous model, CRU Ca release flux was fixed to certain *I*_*spark*_ value (another independent model variable). In the new model it is not fixed anymore, but proportional to difference in [Ca] inside and outside the SR.
3. In the new model Ca release is terminated when it reaches a small value that is comparable with the release amplitude of a single RyR, so that the CICR within the CRU would not sustain any longer in line with modern concepts of spark termination such as induction decay (Laver et al., 2013), spark “death” (Stern et al., 2013), and Ising formalism (Maltsev et al., 2017a).
4. The new model features a more realistic Ca distribution within the cell (Fig. 1). Our previous model was a 2D model: CRUs resided in a square lattice under the cell surface with common pools of SR and cytosol; local Ca concentrations (including LCRs) were predicted by the model only under cell membrane in 2D. The new model includes 3 layers of intracellular voxels. Thus, it becomes a 3D model, making cytoplasmic Ca dynamics more realistic and precise. Its structure is similar to that of Stern model, but the number of voxel layers from cell surface to cell center is limited to three (for more computational efficiency): fine submembrane voxels, intermediate “ring” voxels, and larger cell core voxels. Importantly, the voxel layers are introduced only for computational efficiency and they do not have any physical barriers or gradients. The submembrane voxels (i.e. submembrane space) are not artificially separated from the rest of the cell. All intracellular voxels, including those of submembrane space, have the same characteristics of Ca diffusion and buffering.
5. We introduced junctional SR connected to free SR via a diffusional resistance that determines the kinetics of JSR refilling with Ca that is a key part of the Ca clock mechanism.
6. Ca in junctional SR is buffered by calsequestrin.
7. The CRUs in the prior model positioned as a square lattice. Here we added the capability to introduce various degree of disorder around the ideal square lattice positions of the CRUs so that the resulting distribution of nearest neighbor distances would be spread, i.e. more closely reproduce that reported in confocal microscopy studies (Stern et al., 2014).

**Figure 1.**
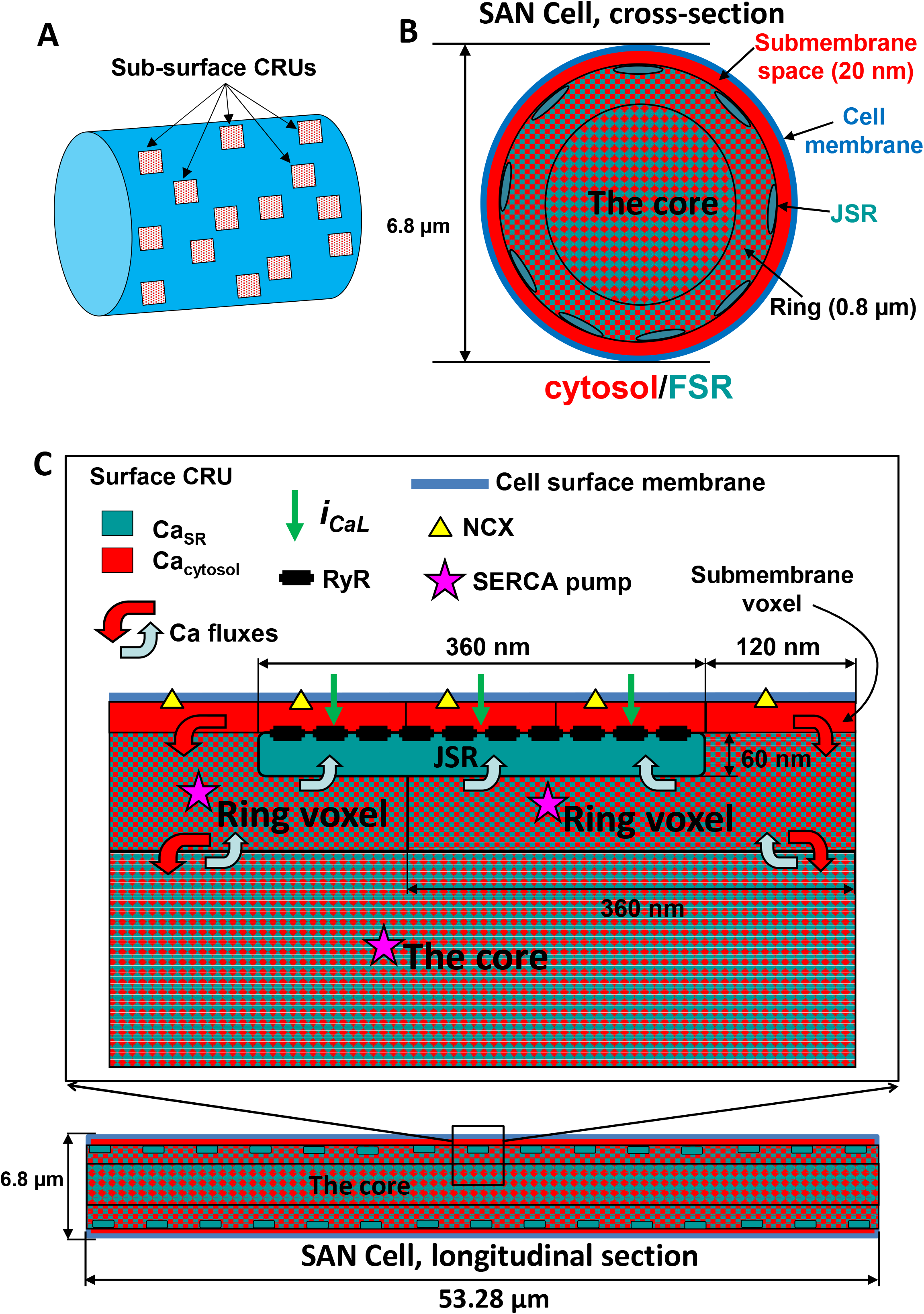
Schematic illustration of approximation of local Ca dynamics in our updated CRU-based SAN cell model: CRUs are placed under the cell membrane (A). Three layers of voxels approximate intracellular Ca dynamics (shown not in scale): in cross-section (B) and longitudinal section (C). For more details see Appendix.

## Results

### The basal AP firing rate increases as CRU distribution becomes disordered

First, we tested how SAN cell function changes if the same number of identical CRUs is distributed differently under cell membrane. We generated and tested cell models with 5 different types of spatial distributions of CRUs with gradually increasing disorder in the CRU positions (Fig. 2, left panels):

1. CRUs placed exactly at the nodes of a square lattice of 1.44 μm size.
2. CRUs slightly deviating from the “square lattice” positions, following Gaussian distribution with standard deviation, SD= 0.25 μm.
3. CRUs moderately deviating from “square lattice” positions, following Gaussian distribution with SD= 0.5 μm.
4. CRUs strongly deviating from the “square lattice” positions, following Gaussian distribution with SD=0.75 μm.
5. Uniformly random CRU positions excluding overlap.

**Figure 2.**
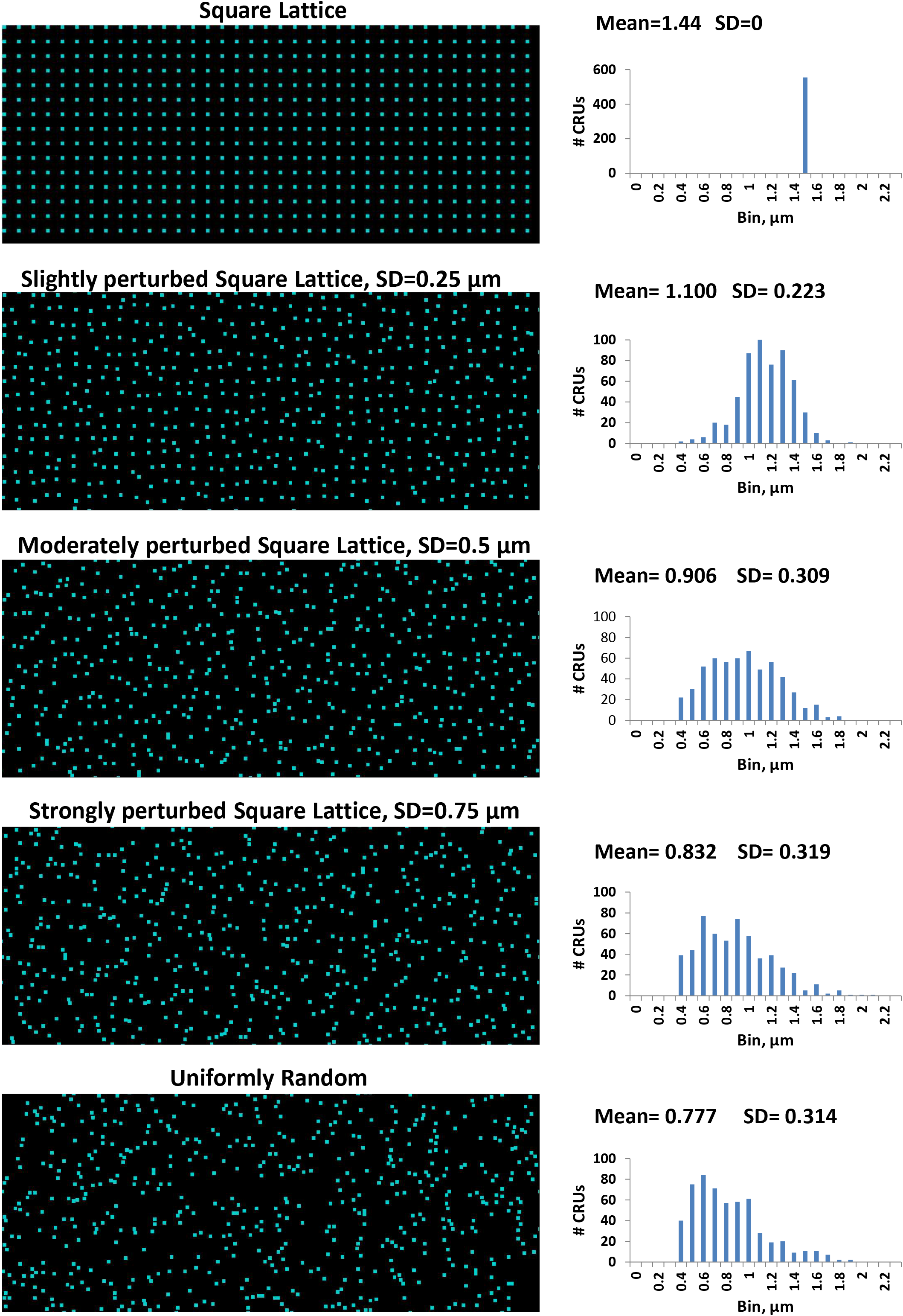
Effect of Poisson clumping: Clusterization of CRUs and emergence of voids as disorder in CRU locations increases. Left panels: Examples of distributions of RyR clusters under the cell membrane with varying degree of disorder used in our simulations of SAN cell function. 555 CRUs are distributed in each case. SD values of normal distributions used to perturb CRU locations from their perfect square lattice positions are shown in respective labels above each image. Right panels: Histograms of the respective distributions of nearest neighbor distances with their mean and SD values (in μm).

As disorder in CRU position increased, the standard deviation of CRU nearest neighbor distance distribution increased, but the mean values decreased from 1.44 μm to 0.777 μm, respectively (Fig. 2, right panels). For a comparison, confocal microscopy measurements performed previously in 6 rabbit SAN cells reported average nearest distances between CRUs in each cell ranging from 0.71 to 0.89 μm (Stern et al., 2014), i.e. close to our simulated values. In all these CRU settings our numerical model simulations generated rhythmic spontaneous AP firing. However, the rate of the spontaneous firing substantially increased as disorder degree increased and average AP cycle length shortened (Fig. 3, for specific values see Table 1, “Basal state” row). The effect of randomness is indeed remarkable: in the extreme case of uniformly random distribution the diastolic depolarization duration was shortened about in half (Fig. 4A, red vs. black traces), and the AP cycle length reduced by >30%, accordingly. This effect is comparable or even exceeds the effect of βAR stimulation on AP cycle length reported in rabbit SAN cells (Vinogradova et al., 2002). The average AP cycle length exhibits a linear dependence vs. average nearest neighbor distance (R^2^=0.96, inset to Fig. 3 A), indicating importance of cell interactions with their nearest neighbors (i.e. CICR) in this phenomenon.

**Table 1.**
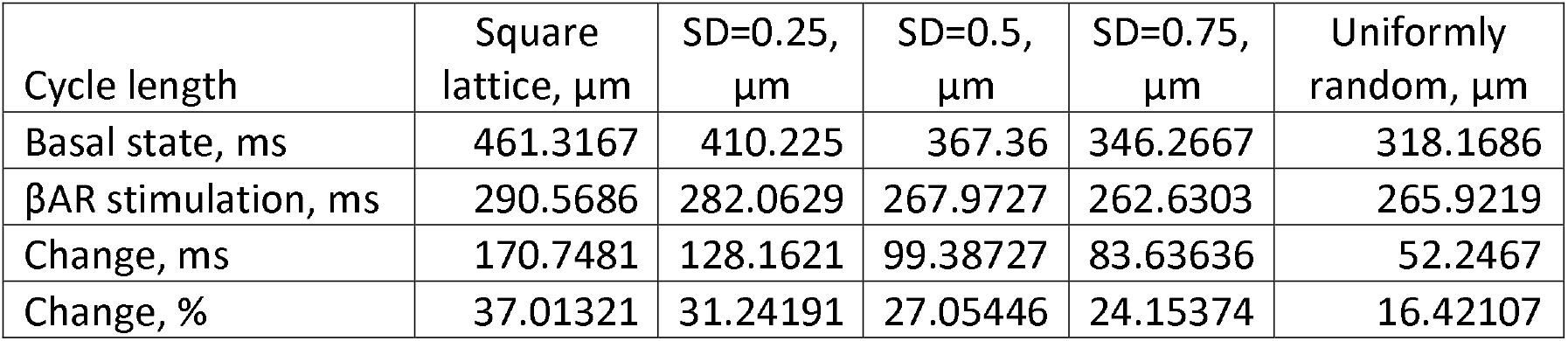
Average values for spontaneous AP cycle length in numerical models with different CRU distributions in basal state and in response to βAR stimulation. AP cycle length was measured for the time interval from 5 s to 16.5 s (when simulations ended). All model simulations began with identical initial conditions and the initial 5s period was omitted from the analysis to allow the system to reach a balance (see our original intervalograms in Fig. A3).

| Cycle length | Square lattice, $\mu\text{m}$ | SD=0.25, $\mu\text{m}$ | SD=0.5, $\mu\text{m}$ | SD=0.75, $\mu\text{m}$ | Uniformly random, $\mu\text{m}$ |
| --- | --- | --- | --- | --- | --- |
| Basal state, ms | 461.3167 | 410.225 | 367.36 | 346.2667 | 318.1686 |
| $\beta\text{AR}$ stimulation, ms | 290.5686 | 282.0629 | 267.9727 | 262.6303 | 265.9219 |
| Change, ms | 170.7481 | 128.1621 | 99.38727 | 83.63636 | 52.2467 |
| Change, % | 37.01321 | 31.24191 | 27.05446 | 24.15374 | 16.42107 |

**Figure 3.**
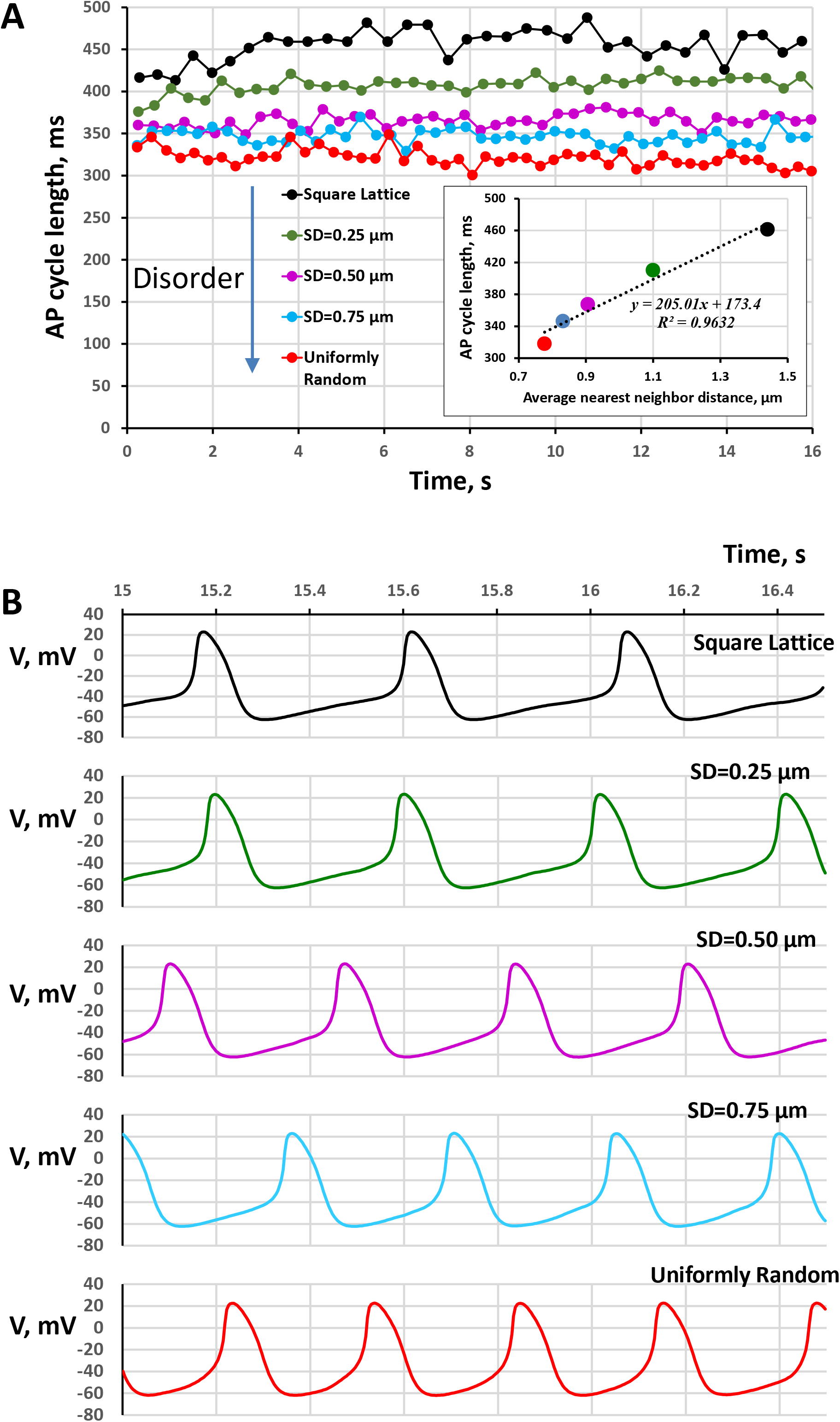
Disorder in CRU position shortens AP cycle length (i.e. increases AP firing rate). A: Intervalogram for 16 s of simulation time for SAN cells with various degrees of disorder in CRU positions. Inset shows that AP cycle length correlates with the average nearest neighbor distance, that is shortens as disorder increases. B: Respective examples of simulated spontaneous APs.

**Figure 4.**
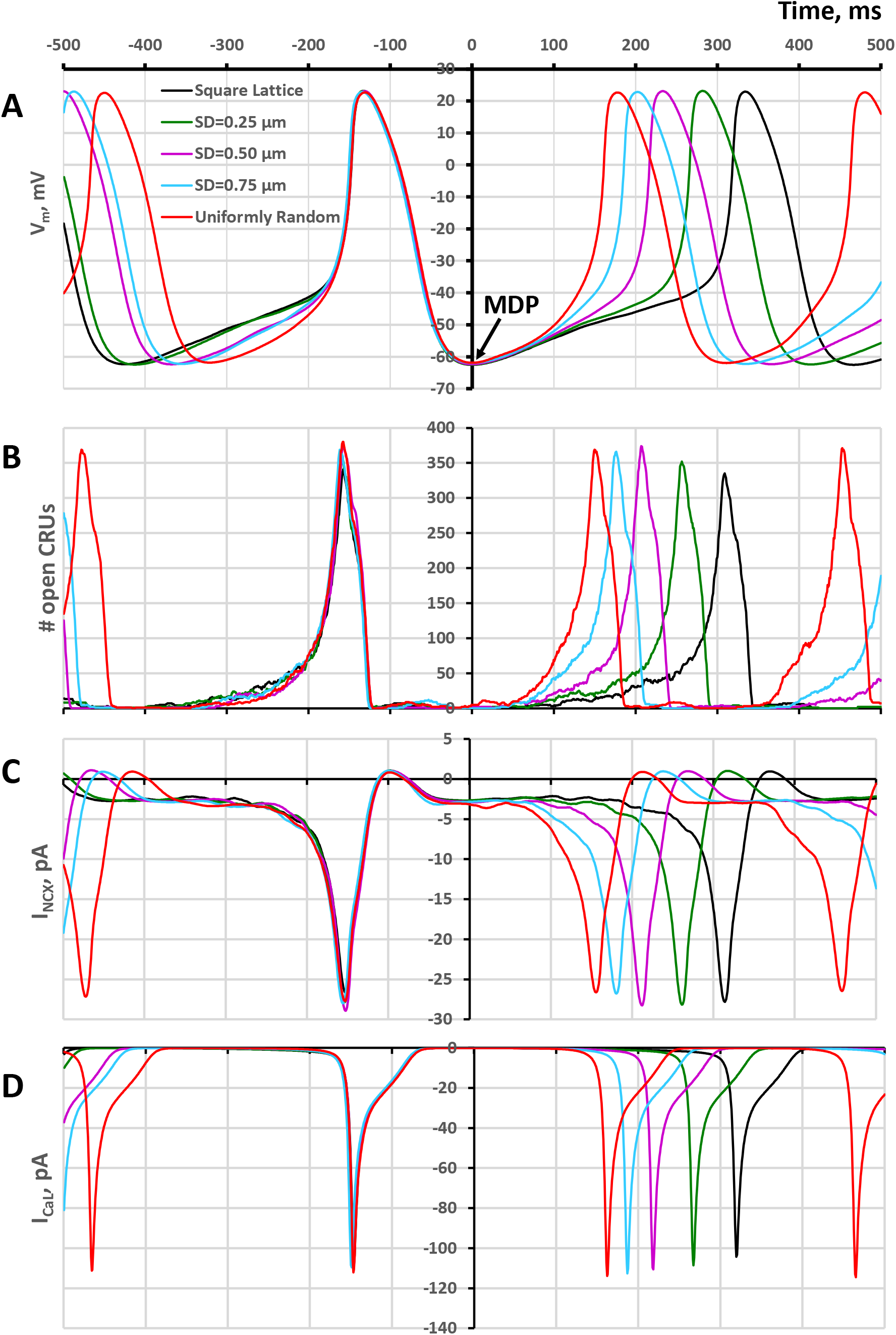
Perturbing CRU positions around their original square lattice positions increases synchronization and recruitment of CRU to release Ca via locally propagating CICR, resulting in earlier activation of *I*_*NCX*_ and shorter AP firing cycle. Shown are representative examples of simulations of *V*_*m*_ (A), # of open CRUs at a given time (B), *I*_*NCX*_ (C), and *I*_*CaL*_ (D).

### The disorder in CRU positions accelerates CRU recruitment and Ca release synchronization in diastole

This is a surprising result, indicating that spatial CRU distribution plays a major role in SAN function. We varied only the locations of CRU, but all other biophysical parameters of the system remained unchanged, i.e. the entire cell electrophysiology, the number and size of CRUs, and Ca cycling parameters including SR Ca pumping. Because our hypothesis was that spatial CRU distribution affects CICR among the CRUs, we examined the dynamics of CRU recruitment in our five RyR settings by monitoring *N*_*CRU*_, i.e. the number of CRU firing at each moment of time during diastole. To compare dynamics of CRU recruitment during diastolic depolarization, we overlapped the traces of representative cycles for each CRU setting and synchronized them at the maximum diastolic potential (MDP) (Fig. 4 A, B). We found that for all CRU settings, a major fraction of CRU pool became recruited to fire during diastolic depolarization, i.e. before the AP upstroke. However, uniform random CRU distribution resulted in a more synchronized and much faster recruitment of CRU firing compared to square lattice setting, with increasing degree of disorder yielding an increase of recruitment pace.

### Electrophysiological consequences of accelerated CRU recruitment: role of *I*_*NCX*_ and *I*_*CaL*_

As a result of faster CRU recruitment, the *I*_*NCX*_ and *I*_*CaL*_ were activated earlier in the cycle, accelerating diastolic depolarization and thereby shortening the AP cycle length (Fig. 4 C,D). In all scenarios *I*_*CaL*_ was activated concurrently with *I*_*NCX*_ or after a short delay (Fig. 5), indicating that the system operates via the coupled-clock paradigm (Maltsev and Lakatta, 2009) and in accord with ignition theory (Lyashkov et al., 2018). The short delay increased with the increasing disorder in CRU positions, indicating that enhanced CRU recruitment activates *I*_*NCX*_ at low voltages, below *I*_*CaL*_ activation (<-50 mV). To illustrate the diastolic CRU recruitment, we made representative screenshots of CRU firing for each CRU setting overlapped with instant local Ca distribution under the cell membrane at the membrane potential (*V*_*m*_) of −45 mV that is close to the AP ignition onset and *I*_*CaL*_ activation threshold (Fig. 6). Mainly individual sparks were observed in the square lattice setting. In contrast, uniform random setting showed propagating LCRs. The intermediate setting with various degrees of disorder exhibited intermediate recruitment. Different CRU firing patterns can also be seen in the respective movies of local Ca dynamics in subspace (Movies 1-5).

**Figure 5.**
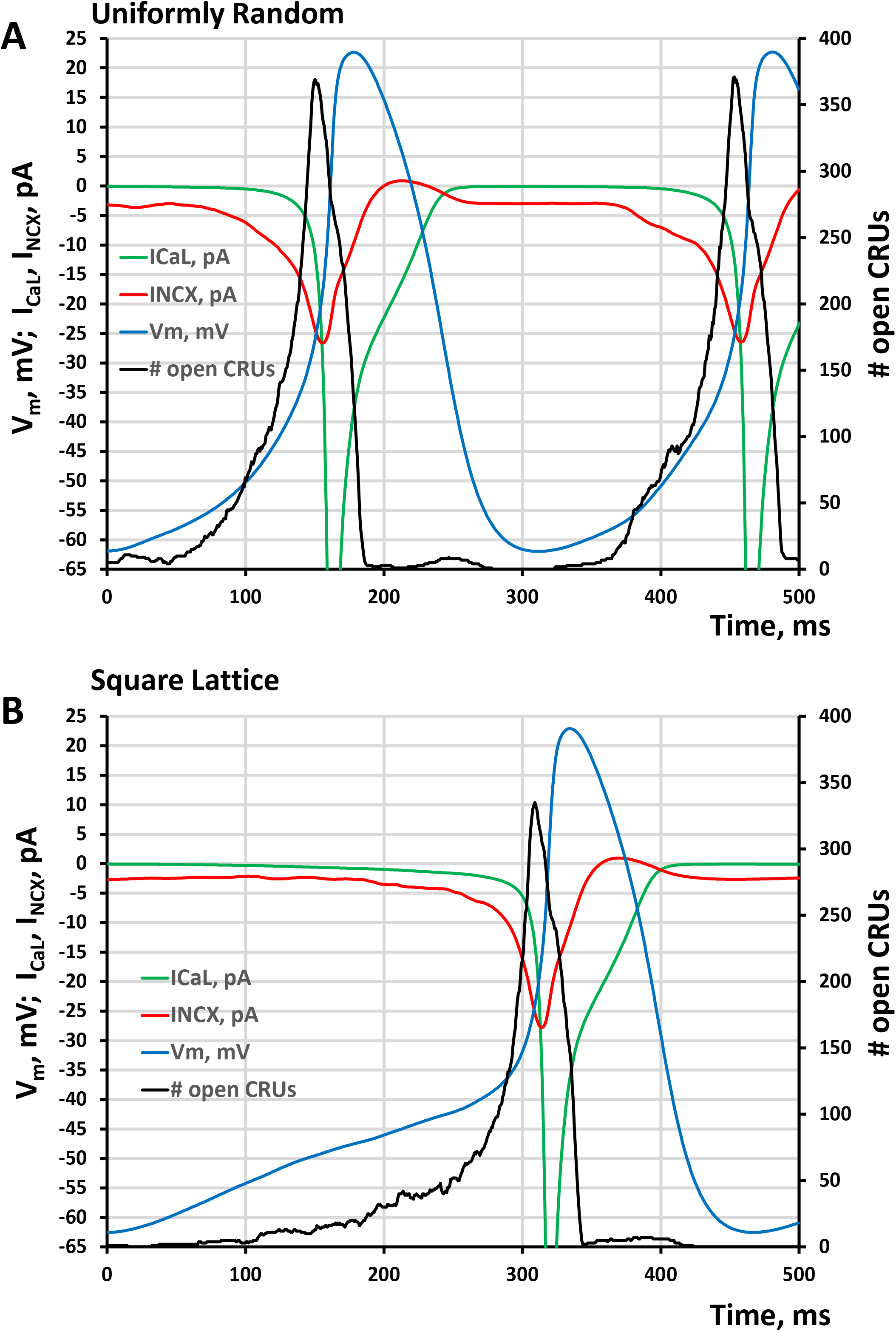
Disorder in CRU positions accelerates synchronization of CRU firing and AP ignition process. Shown are representative cycles of AP firing (*V*_*m*_, blue curves) for 500 ms simulation time beginning from the maximum diastolic polarization (MDP, time=0) for two CRU distributions: Uniformly random (A) and square lattice (B). The ignition process in both cases is characterized by simultaneous activation of CRUs firing (black), *I*_*NCX*_ (red), and *I*_*CaL*_ (green, truncated), which *V*_*m*_ to the AP activation threshold. However, the activation kinetics are faster in the uniformly random case so its diastolic depolarization is about twice shorter.

**Figure 6.**
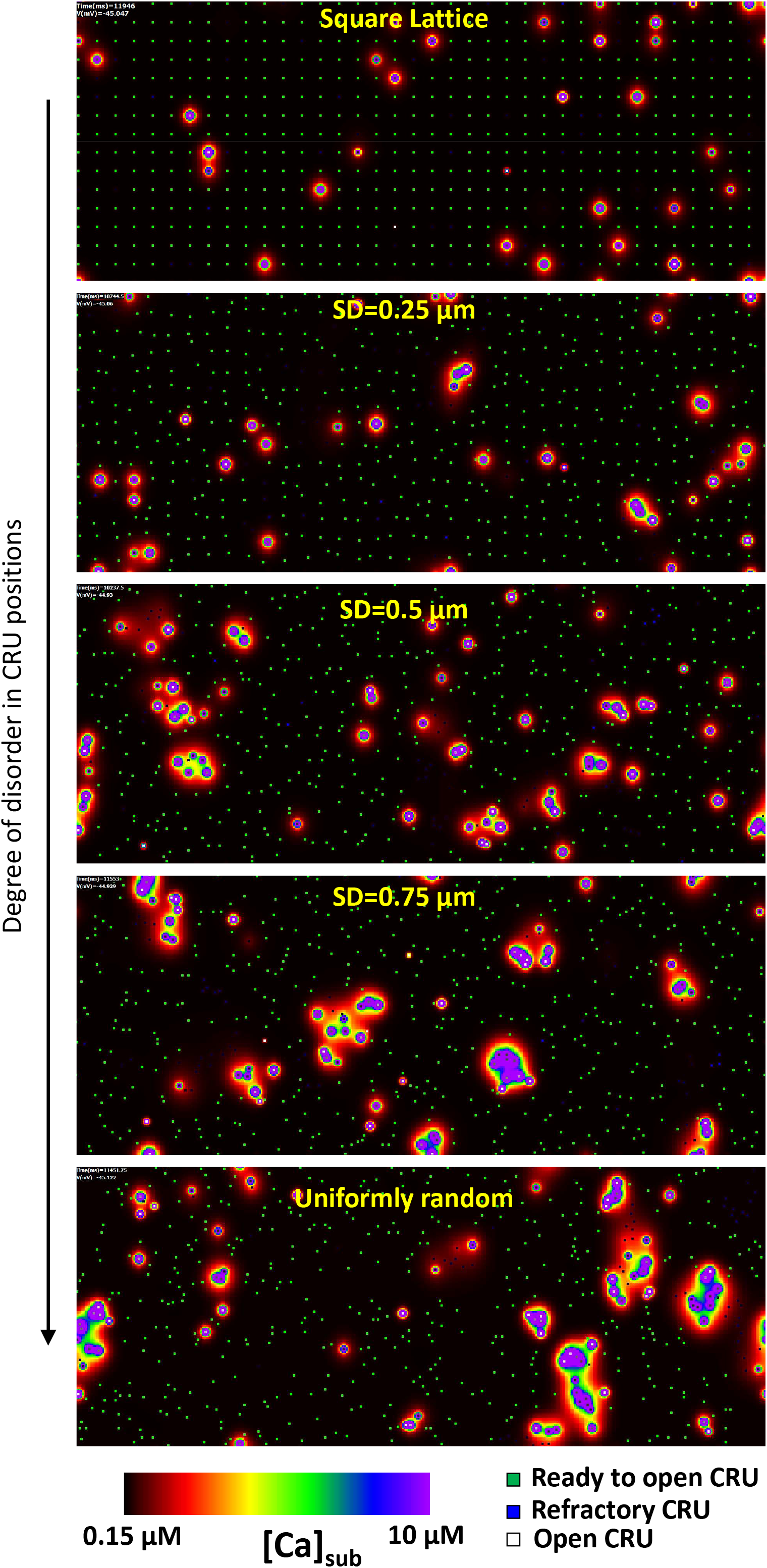
Sizes of LCRs increase during diastolic depolarization via propagating CICR as disorder in CRU positions increases from square lattice to uniformly random (from top to bottom). Shown are distributions of local *[Ca]*_*sub*_ ([Ca] under cell membrane) at the same membrane potential of −45 mV, i.e. close to the *I*_*CaL*_ activation threshold. [Ca]_sub_ is coded by a color scheme shown at the bottom from 0.15 μM to 10 μM; open CRUs are shown by white dots. Closed CRUs in refractory period are shown by blue dots; closed reactivated CRUs (available to fire) are shown by green dots. JSR Ca level is coded by respective shade (white, blue or green) with a saturation level set at 0.3 mM. See respective Movies 1-5.

### Disorder in CRU positions increases robustness of basal automaticity

SAN cells are very heterogeneous in their biophysical properties and their key ion current densities vary substantially, e.g. up to an order of magnitude for *I*_*CaL*_ (Honjo et al., 1996; Monfredi et al., 2017) (Fig.7, red circles). On the other hand, our simulations showed that the rate of spontaneous AP firing can be substantially reduced because of differences in spatial CRU distribution (Fig. 3). We then hypothesized that it is possible to find a range for realistic *I*_*CaL*_ densities at which spontaneous AP firing is impossible for the square lattice CRU distribution, but possible for uniformly random model. To match realistic *I*_*CaL*_ density values (in pA/pF) and our model parameter *g*_*CaL*_ (in nS/pF) determining maximum *I*_*CaL*_ conductance, we performed voltage-clamp simulations (Fig. 7 B) with the voltage step protocol similar to that in experimental studies (Honjo et al., 1996; Monfredi et al., 2017). To instantly attain steady-state activation and inactivation gating, we set respective gating variables *fL* and *dL* in the model to their steady-state values calculated at the holding potential of −45 mV. The I/V curve for peak values of simulated voltage-clamp *I*_*CaL*_ traces are shown in the inset. Thus, our basal state conductance 0.464 nS/pA in the model generated the maximum peak *I*_*CaL*_ current of 11.17 pA/pF shown by upper edge of the magenta band in Fig. 7 A that crosses the range of the experimentally measured *I*_*CaL*_ densities approximately in the middle. Note: the graph in Fig. 7 A has a dual *y* axis scale, for *I*_*CaL*_ and *g*_*CaL*_ respectively, bridging experimental and simulated data.

**Figure 7.**
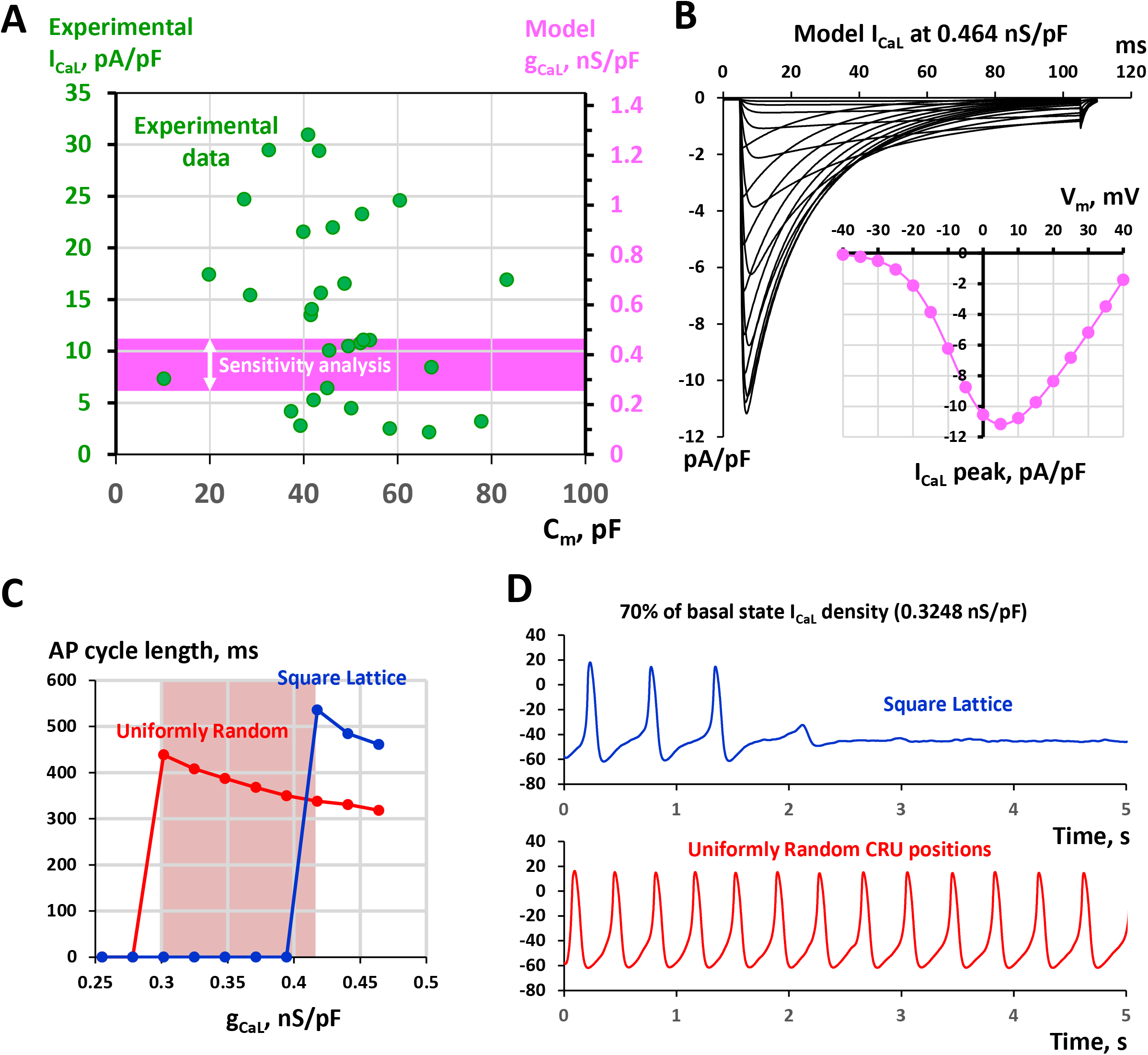
Results of sensitivity analysis demonstrating failure of pacemaker function (AP firing ceased) in cell models with square lattice positions of CRUs, but not with uniformly random CRU distribution at lower, albeit physiological, *I*_*CaL*_ densities. A: Bridging experimentally measured *I*_*CaL*_ densities (green circles) and our model parameter *g*_*CaL*_ describing maximum *I*_*CaL*_ conductance (magenta). The data points for *I*_*CaL*_ experimental data were replotted from (Monfredi et al., 2017). Magenta band shows range of *g*_*CaL*_ in our sensitivity analysis. B: Voltage clamp simulations of *I*_*CaL*_ traces (black) to reproduce experimental protocol and I/V relationship (inset) to find the maximum peak of *I*_*CaL*_ in the model to match respective *I*_*CaL*_ densities measured experimentally (upper edge of the magenta band in panel A). C: Results of sensitivity analyses for cell models with uniformly random CRU distribution (red) and square lattice distribution (blue). Red shade square shows the *g*_*CaL*_ margin where square lattice models failed, but uniformly random model continued generating spontaneous APs. D: An example of AP firing failure in a square lattice model, but rhythmic AP firing in uniformly random model when in both cases *g*_*CaL*_ reduced to 70% of its basal value. See original simulation data in Fig. A2

Then we performed a wide range sensitivity analysis for *g*_*CaL*_ from its basal value of 0.464 nS/pF (100%) down to 0.2552 nS/pF (55%) that remained within the range of realistic *I*_*CaL*_ values (magenta band in Fig. 7 A). The analysis revealed a wide margin of *g*_*CaL*_ values (shown by red shade in Fig. 7 C) where the AP firing is impossible with square lattice distributions of CRUs (red circles), but possible with uniformly random CRU positions. All simulated traces of 16.5 s duration are shown in Fig. A2 and an example of the result is shown in Fig. 7 D for *g*_*CaL*_ of 0.3248 nS/pF.

This analysis revealed another important aspect of SAN function with different CRU distributions: the model with the square lattice CRU distribution was capable of generating AP firing with relatively long cycle length of 536 ms on average (before it failed), whereas in uniformly random CRU model could reach only 438 ms. Thus, SAN cells can (in theory) harness their CRU distribution to safely reach low rates.

### Subthreshold signaling

A newly discovered paradigm pacemaker cell function is the ability of some SAN cells to reversibly switch to dormant (non-firing) state that was shown in isolated cells (Kim et al., 2018; Tsutsui et al., 2018; Tsutsui et al., 2021) and in SAN tissue (Bychkov et al., 2020; Fenske et al., 2020). Furthermore, it was shown that non-firing cells in SAN tissue can generate subthreshold signals that were proposed to be important signaling events for generation of synchronized cardiac impulses (Bychkov et al., 2020). Thus, we hypothesized that CRU distribution is important not only for AP firing, but also for generation of the subthreshold signals in dormant cells. One important factor in cell dormancy is a lower density of *I*_*CaL*_ (Tsutsui et al., 2021). Using the results of our sensitivity analysis, we chose a low *g*_*CaL*_ of 0.2552 nS/pF, at which AP firing is impossible in both uniformly random and square lattice models (Fig. 8A) and compared subthreshold *V*_*m*_ signals in the non-firing cells in each case. We found that subthreshold *V*_*m*_ oscillations in case of uniformly random distribution are much more powerful and occurred at a faster frequency vs. those in the square lattice model as evident from the overlapped power spectra of both *V*_*m*_ signals (Fig. 8B) computed for the time period after AP failure.

**Figure 8.**
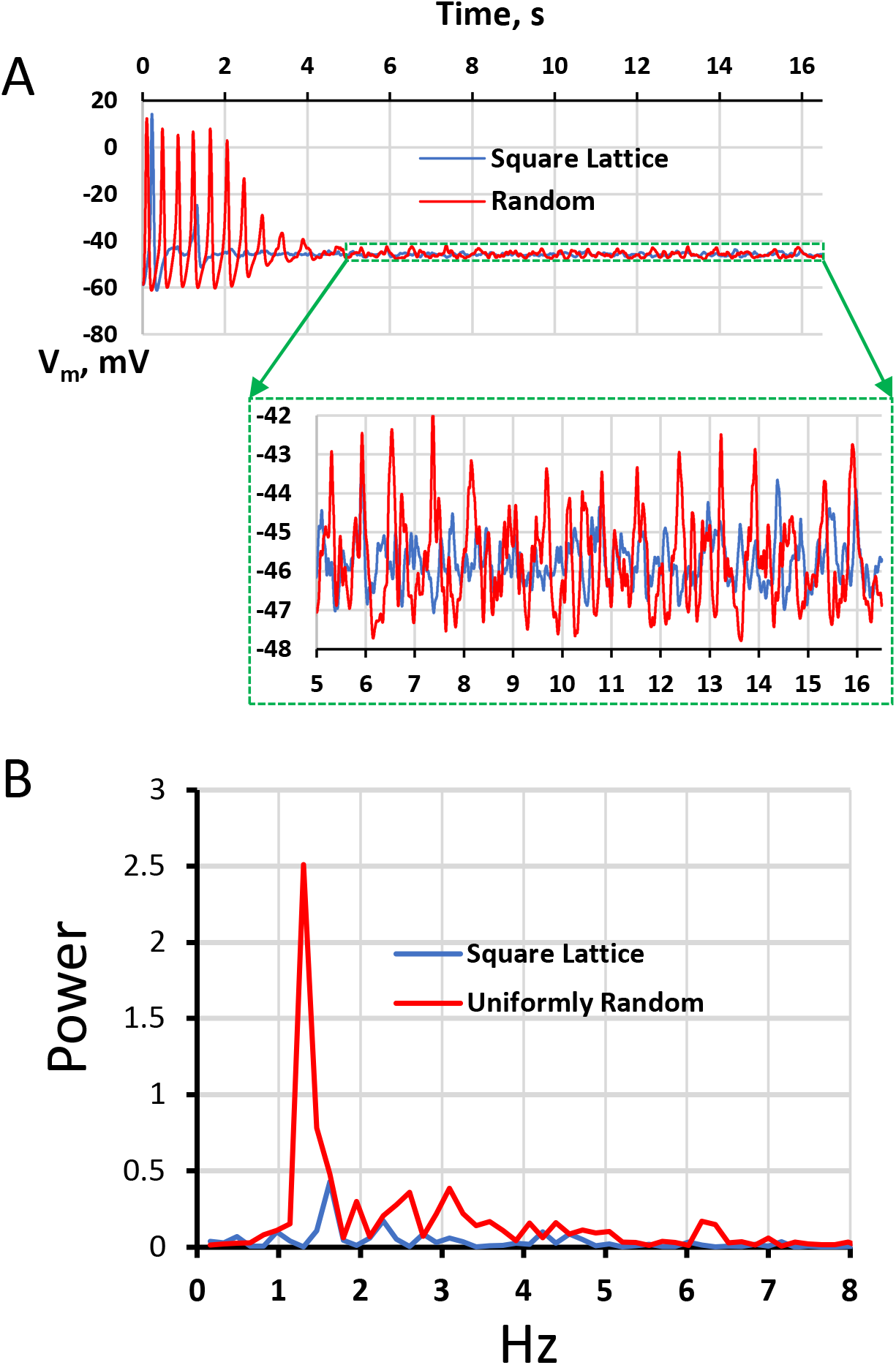
New insights for dormant cell signaling. A: Despite both models with uniformly random and square lattice models failed at a very low *g*_*CaL*_ of 0.2552 nS/pF, subthreshold *V*_*m*_ oscillations continue in both cases. However, oscillations in uniformly random distribution had much larger amplitude (shown in inset). B: The oscillations were not only much more powerful but occurred at a faster frequency as revealed by the power spectra of both *V*_*m*_ signals computed for time after AP failure (from 0.5 s to 16.5 s).

### Order in CRU position increases chronotropic reserve in fight-or-flight reflex

Normal pacemaker cell function is not only robust, but also flexible, i.e. able to increase the AP firing rate under stress (fight-or-flight reflex) or decrease it at rest. We tested if CRU distribution plays a role in the flexibility of SAN cell function by examining responses of our model to βAR stimulation. Our simulations showed that all five CRU settings from square lattice to uniform random exhibit fight-or-flight reflex, i.e. the AP cycle length notably decreased in the presence of βAR stimulation (red bars vs. blue bars in Fig. 9A). All intervalograms are shown in Fig. A3, and examples of βAR stimulation effect in extreme cases of CRU distributions are shown in Fig. 9B. The presence of disorder substantially decreased the stimulation effect, i.e. the AP cycle length shortening decreased as disorder in CRU positions increased (both absolute and relative changes are given in Table 1). Furthermore, the absolute decrease in the AP cycle length was linearly related (R^2^=0.9931) to the basal cycle length before βAR stimulation (Fig. 9C) that is in accord with a recent experimental report (Kim et al., 2021).

**Figure 9.**
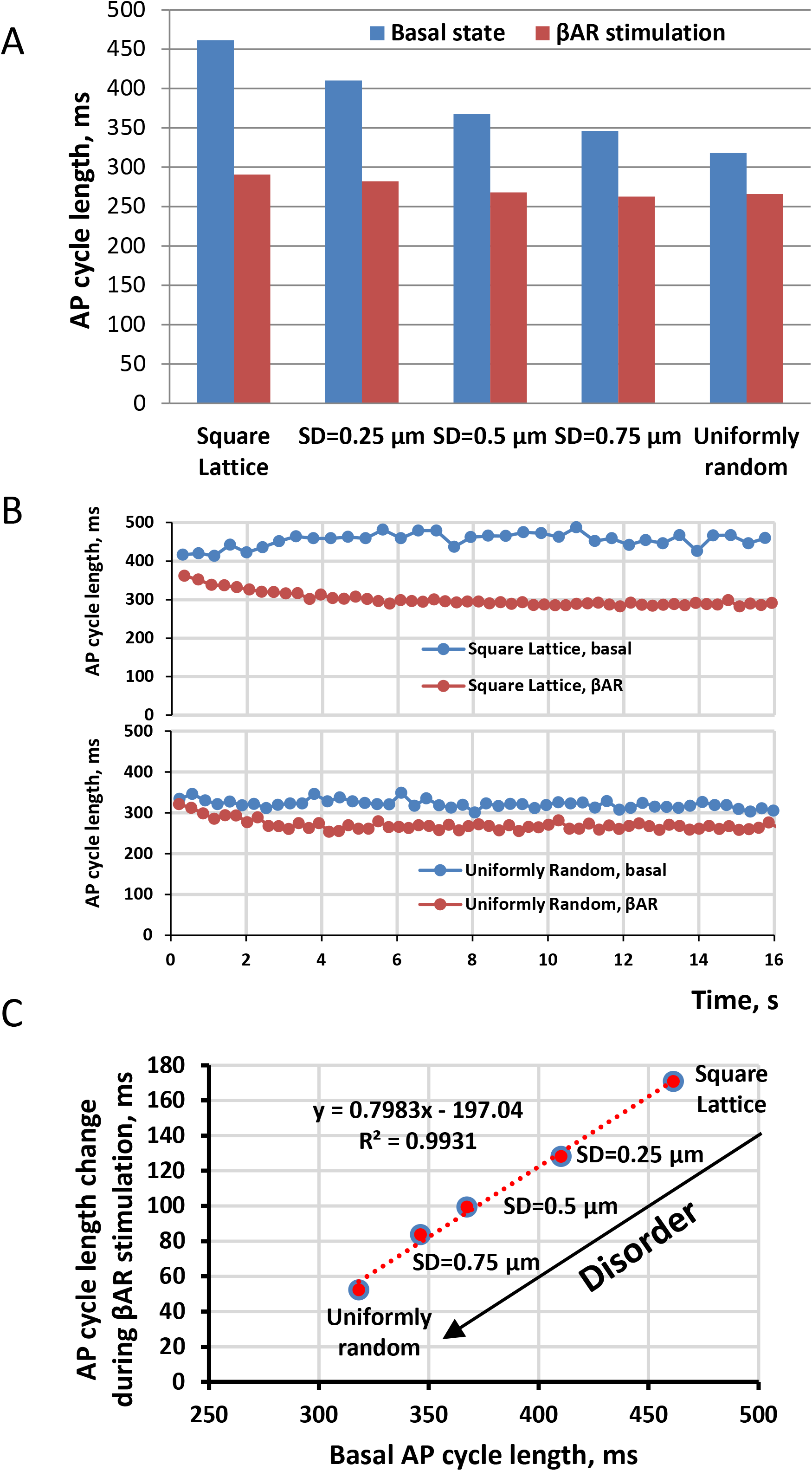
Disorder in CRU positions shortens the AP cycle length, but simultaneously decreases the effect of β-adrenergic receptor stimulation. A: Bar graph of respective AP firing lengths (for numerical data, see Table 1). B: An example of simulations of βAR stimulation effects for extreme cases of CRU distributions: uniformly random (upper panel) vs. square lattice (bottom panel). C: AP cycle length changes correlate with basal AP cycle length (before βAR stimulation), βAR stimulation synchronizes AP firing towards a common AP firing cycle length. See original simulation data in Fig. A3

To get further insights into the effect of spatial disorder on βAR stimulation, we simulated and compared the dynamics of *N*_*CRU*_ and *I*_*NCX*_ during a representative cycle for square lattice and uniform models (Fig. 10). We found that in both cases the βAR stimulation effect, linked to earlier and stronger recruitment of CRUs, translated to respective earlier and stronger activation of *I*_*NCX*_. However, this effect (i.e. the shift to earlier CRU recruitment) was stronger in square lattice setting, indicating the presence of a larger “reserve” CRU pool ready to be activated and synchronized via βAR stimulation.

**Figure 10.**
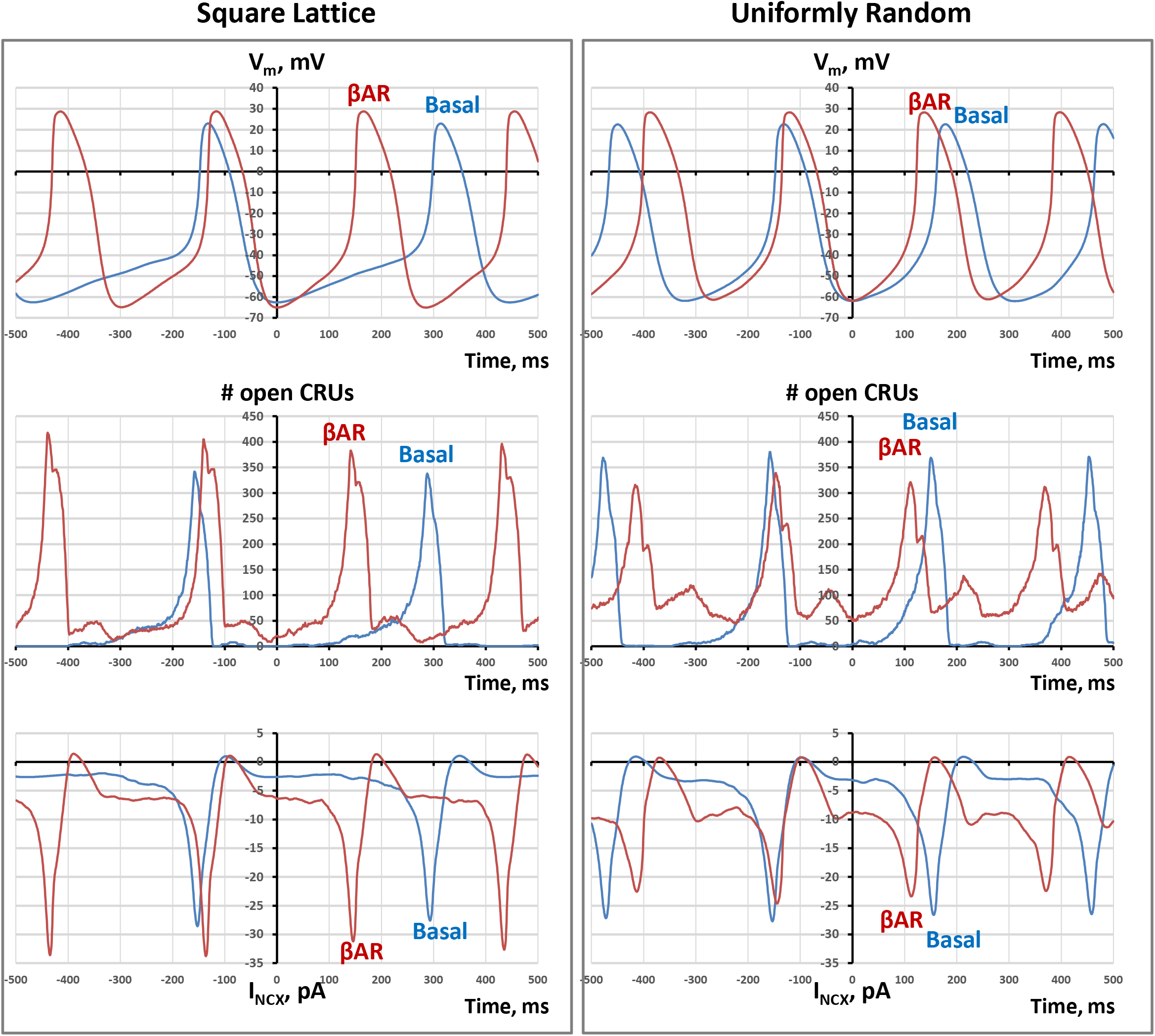
Numerical simulations illustrating the mechanism of a stronger reduction of AP cycle length by βAR stimulation (βARs) in square lattice vs. uniform random CRU spatial distribution (*V*_*m*_, top pannels). The stronger effect in the lattice case is due to its stronger effect on CRU recruitment (*N*_*CRU*_, middle panels) and its attendant activation of *I*_*NCX*_ (bottom panels). Disorder in CRU positions substantially shortens the AP cycle length in basal state, but simultaneously decreases the effect of stimulation.

## Discussion

### Results summary

The focus of the present study was to investigate if disorder in CRU positions under cell membrane has any notable effect on pacemaker SAN function. We used an upgraded numerical SAN cell model featuring Ca clock at the level of local CRU network, coupled to the cell membrane electrophysiology equations. As the extent of disorder in the CRU positions increased, SAN function was assessed via its spontaneous AP firing rate or cycle length. We obtained an intriguing result: spatial disorder actually increased both the firing rate and robustness of pacemaker function in the basal state. The disorder decreased the CRU nearest neighbor distances and facilitated Ca release propagation via CICR, leading to earlier and stronger LCR signal that increased AP firing rate and also initiated AP firing in cells that could not fire APs with fully ordered square lattice positioning of CRU. The magnitude of the effect on the firing rate was substantial, quantitatively similar to that known for βAR stimulation. However, the boost of robustness bears a cost in the flexibility of pacemaker function. The range of AP firing rate modulation by βAR stimulation substantially narrowed as disorder and its attendant robustness increase. This happens because the disorder-facilitated release synchronization is utilized a majority of CRUs in the basal state and provides a smaller contribution to rate increase during stimulation due to fewer unutilized “reserve” CRUs.

### Upgraded CRU-based model of SAN cell

An important result of the present study was a major upgrade of our previous CRU-based model of rabbit SAN cell (Maltsev et al., 2011; Maltsev et al., 2013) (Fig.1, see Appendix for more details). Timing for CRU activation was previously defined phenomenologically via a fixed refractory period given as an independent variable followed by a Poisson process, whereas termination was set simply to a mean value of spark duration. Both the refractory period and spark duration were taken directly from experimental measurements. The new model includes Ca release activation and termination mechanisms linked SR Ca load as shown in experimental and theoretical studies (Zima et al., 2008; Maltsev and Lakatta, 2009; Imtiaz et al., 2010; Vinogradova et al., 2010; Zima et al., 2010; Laver et al., 2013; Stern et al., 2013; Maltsev et al., 2017a; Veron et al., 2021). These model enhancements are of crucial importance for this and future studies. Firstly, the new model reflects recent progress in our understanding of RyR function. Secondly, it describes mechanistically the coupling of the Ca clock to the membrane clock at the scale of the local CRU network. This is an important niche in the variety of numerical SAN cell models developed thus far. As described in details in Methods section and Appendix, its unique scale is positioned between simple common pool models (e.g. (Kurata et al., 2002; Maltsev and Lakatta, 2009),(Severi et al., 2012)) and extremely complex RyR-based models (Stern et al., 2014; Maltsev et al., 2017b).

The models at each level of signal integration are important to understanding the “big picture”, i.e. how the phenomenon of heartbeat emerges, bridging the gaps between scales (Clancy and Santana, 2020; Weiss and Qu, 2020) in the spirit of multiscale modeling (Qu et al., 2011). The CRU-based modeling of SAN cells has become especially important with respect to recent discoveries of importance of local Ca signaling for SAN tissue function (Bychkov et al., 2020). It was shown that Ca signals are markedly heterogeneous in space, amplitude, frequency, and phase among cells comprising an HCN4+/CX43- cell meshwork of SAN, and synchronized APs emerge from heterogeneous subcellular subthreshold Ca signals (modelled here, see Fig. 8)). While a series of insightful multicellular models of SAN tissue have been developed (see for example (Oren and Clancy, 2010; Inada et al., 2014; Gratz et al., 2018; Li et al., 2018)), local Ca signaling has not been numerically studied at the tissue level yet. Our new CRU-based model generates at a low computational cost LCRs observed experimentally both in isolated cell and in intact SAN and, thus, can be used as a functional unit in future multi-cellular modeling to investigate the role of local Ca signaling at the level of SAN tissue that represents the frontier of heart pacemaker research (Clancy and Santana, 2020; Weiss and Qu, 2020).

### Mechanisms of the CRU spatial disorder effect

Empty spaces and clusters are an intrinsic feature of random spatial distributions, known as Poisson clumping. Such emerging CRU clumps are clearly seen by eye in our examples of 2D representations of CRU networks in Fig. 2 (left panels), and the clustering effect is quantitatively manifested by a broader distribution of nearest neighbor distances with notably shorter averages (right panels). Thus, the disorder in CRU positions creates shortcuts and opportunities for CICR propagation and thus CRU recruitment and synchronization. Once the Ca release becomes more synchronized, LCR sizes increase, the amplitude of LCR net diastolic signal also increases, and, very importantly, the timing of net LCR signal simultaneously shortens, as we previously demonstrated (Maltsev et al., 2011). Further effect of LCRs on AP firing rate is executed via NCX and *I*_*CaL*_ to accelerate diastolic depolarization (Figs 4,5, and 10) as postulated in the coupled-clock theory (Maltsev and Lakatta, 2009; Lakatta et al., 2010) and more recent ignition theory (Lyashkov et al., 2018), in line with previous numerical studies in a CRU-based SAN cell model (Maltsev et al., 2013) and RyR-based model (Stern et al., 2014).

The Ca release synchronization mechanism via local CRU recruitment is impacted by many factors including for example *I*_*spark*_, i.e. Ca release flux of a single CRU which we studied previously (Maltsev et al., 2011). It is determined by SR Ca refilling kinetics, driven by PKA-dependent phosphorylation of Ca cycling proteins (Vinogradova et al., 2006; Lakatta et al., 2010). Mechanistically speaking, Ca clock ticks during diastole when SR refills to a certain threshold level so that Ca current via a single RyR channel becomes big enough to recruit its neighboring RyRs within CRU to generate a spark (Zima et al., 2010; Veron et al., 2021) and then *I*_*spark*_ becomes large enough to generate an LCR, i.e. a series of propagating sparks that depends on the local CRU distribution. Thus, the recruitment phase becomes delayed if the nearest neighbor distances are larger (such as in square lattice arrangement) and require a larger SR Ca loading commensurate with larger *I*_*spark*_ to begin with. And vice versa, the recruitment starts earlier in the cycle if the nearest neighbor distances are shorter (such as in uniformly random arrangement).

With larger nearest neighbor distances in the square lattice setting many CRUs never fire during diastole in basal state beating, but become recruited during βAR stimulation (Fig. 10). For example, in basal state on average about 44% of CRUs fired at −35 mV (i.e. the end of diastolic depolarization) in the uniformly random model, but only 29% in the lattice model; however, during βAR stimulation about 53% of CRUs fired in either case, i.e. CRUs became equally well recruited independent of the model. Thus, the slower recruitment and the non-recruited CRUs in the basal firing represent chronotropic reserve mechanisms that are utilized during βAR stimulation. The reserve is obviously limited by the number of CRUs, and if more CRUs are recruited to fire in each diastole in basal state, then the number of CRUs in reserve shrinks (and vice versa); that is why βAR stimulation effect is much smaller in the uniformly random model vs. lattice model. This result is evidence of possible functional importance of the chronotropic reserve linked to CRU recruitment: We have previously demonstrated that CRU recruitment stabilizes diastolic I_NCX_ amplitude that explained, for example, a paradoxical effect of partial knockout of NCX in mice to reduce chronotropic reserve with no effect on the basal rate (Maltsev et al., 2013).

### Possible importance for normal function, pathological conditions, and aging

Thus far CRU distribution in SAN cells has not been systematically studied. However available data indicate that locations of CRUs in SAN cells do not form a perfect grid, but exhibit a notable degree of randomness (Lyashkov et al., 2007; Stern et al., 2014). It has been shown that different SAN cells may have different CRU arrangement: central SAN cells have a substantial degree of disorder in their RyR cluster positions, whereas peripheral cells feature striations and more organized RyR clustering (Rigg et al., 2000; Musa et al., 2002). Our result may indicate that the central cells are capable of robustly generating high frequency signals, but unlikely to have a substantial CRU recruitment reserve to utilize during βAR stimulation (they may still have other mechanisms to increase their rate). On the contrary, the cells with more organized RyR clustering (wherever they are) may have a larger CRU recruitment reserve.

Thus, the real CRU distribution is neither uniformly random, nor perfectly spaced. By adjusting the CRU positioning, SAN cells can reach a balance of robustness and flexibility. This balance is required (and dictated) for each cell by its specific location and functional role within the cellular network of SAN tissue. This could be important for regulation of βAR stimulation response among individual cells within SAN tissue for its optimal integrated chronotropic response (Brennan et al., 2020; Yuan et al., 2020; Kim et al., 2021). Here we show that the AP cycle length decrease during βAR response depends linearly on the cycle length in the basal state. This finding is line with recent experimental results that βAR stimulation synchronizes a broad spectrum of AP firing rates in SAN cells toward a higher population average (Kim et al., 2021).

Another possible importance of the disorder-facilitated AP firing could be regulation of cell dormancy, a recently discovered phenomenon manifested by the absence of automaticity of SAN cells that can be acquired via βAR stimulation (Kim et al., 2018; Tsutsui et al., 2018; Tsutsui et al., 2021). One important factor of cell dormancy is a lower density of *I*_*CaL*_ (Tsutsui et al., 2021). Here we show that by rearranging its CRU positions, a SAN cell with a low *I*_*CaL*_ density can switch its functional state between being dormant or AP firing, as illustrated in Fig. 7. Nonfiring cells have also been recently found in intact, fully functional SA node (Bychkov et al., 2020; Fenske et al., 2020); their numbers substantially varied in different chronotropic states of the SAN (Fenske et al., 2020). A new pacemaker mechanism has been proposed that synchronized cardiac impulses emerge from heterogeneous local Ca signals within and among cells of pacemaker tissue, including subthreshold signals in non-firing cells (Bychkov et al., 2020). Here we show that subthreshold *V*_*m*_ oscillations in case of uniformly random distribution are much more powerful and occurred at a faster frequency vs. those in the square lattice model (Fig. 8). Thus, subthreshold signaling can be also regulated by the spatial CRU distribution.

Another interesting result of our sensitivity analysis is that the model with the perfect square lattice CRU locations (before it failed) was capable of generating AP firing with much longer cycle length (i.e. at a low rate of excitations), whereas in uniformly random scenario the model failed at a shorter cycle length (i.e. cannot reach lower rates) (Fig. 7C). Thus, pacemaker cells can harness this mechanism linked to CRU distribution to reach lower rates in addition to other known bradycardic mechanisms such as shift in voltage activation of I_f_ (DiFrancesco and Tromba, 1988), I_K,Ach_, and protein dephosphorylation (decreasing clock coupling) (Lyashkov et al., 2009).

Pathological conditions and aging are usually associated with disorder in molecular positions, interactions, and functions. We demonstrate that excessive disorder in CRU positions within SAN cells decreases fight-or-flight response while shrinking the range of lower AP rates (Figs 7 C and 9), and hence can explain, in part, the limited range of heart rates associated with age and in disease. On the other hand, the mechanism of disorder-facilitated Ca release propagation can act together with increased sympathetic tone to compensate the age-intrinsic heart rate range decline with age (Tsutsui et al., 2016). With respect to atrial and ventricular cells, excessive disorder in CRU positions (in cardiac disease) is expected to facilitate Ca release propagation, i.e. waves formation, increasing the risk of life-threatening arrhythmia (Ter Keurs and Boyden, 2007).

### A broader interpretation: living systems harness disorder to function

While noise is a broad term that is usually associated with undesirable disturbances or fluctuations, many biological systems harness disorder to function. Randomness creates opportunities to exceed a threshold that is above the mean, analogous to the way a quantum particle can tunnel across a barrier while a classical deterministic particle cannot. It can improve signal transmission or detection, e.g. via stochastic resonance (McDonnell and Abbott, 2009). Random parameter heterogeneity among oscillators can consistently rescue the system from losing synchrony (Zhang et al., 2021). Randomness is critical for cardiac muscle cell function. The local control theory developed by Michael Stern in 1992 (Stern, 1992) predicted the Ca sparks (found later experimentally (Cheng et al., 1993)) and explained smooth regulation of excitation-contraction coupling in cardiac muscle via statistics of success and failure of a CRU to generate a spark when L-type Ca channels open. We have recently shown that statistical physics approach is also helpful to understand spontaneous Ca spark activation and termination (via Ising formalism (Maltsev et al., 2017a; Maltsev et al., 2019; Veron et al., 2021)). In the present study we show that disorder could be also important for cardiac pacemaker function: disorder in CRU locations determines statistics of success and failures for a firing CRU to recruit to fire its neighboring CRU, observed as propagating LCRs that are critical for SAN cell function (Lakatta et al., 2010). Thus, paradoxically disorder in CRU positions facilitates functional order in terms of LCR emergence via self-organization by means of positive feedback provided by CICR, culminating in higher AP firing rates, whereas order in CRU positions is associated with individual stochastic sparks, i.e. functional disorder for a major part of the diastolic depolarization duration, culminating in lower AP firing rates. A broader interpretation of our results is that disorder in a network featuring diffusion-reaction interactions can facilitate excitation propagation, that may be applicable to RyR arrangement within a CRU to generate a spark (down-scale) or cell-to-cell interactions in SAN tissue to generate cardiac impulse (up-scale).

### Limitations and future studies

In our previous study we showed that a CRU network lacking release propagation can acquire release propagation capability by introducing a subset of smaller “bridging” CRUs that create propagation shortcuts and allow sparks to jump from one firing CRU to its neighbor (Stern et al., 2014). In the present study we show that bridging of CRU network is not required to achieve release propagation: the intrinsic disorder in CRU positions can naturally create the bridges and propagation shortcuts without additional bridging CRUs. On the other hand, our study is of a reductionist type focused on the disorder contribution, whereas the realistic CRU distribution in SAN cells has more complex, hierarchical structure that includes CRUs of various sizes (Stern et al., 2014). Thus, future studies will clarify the role of disorder in the more realistic settings with different CRU sizes and more precise CRU locations within the cell measured by super resolution microscopy in 3D (pilot studies (Maltsev et al., 2016; Greiser et al., 2020)) rather than by confocal microscopy in tangential sections (Stern et al., 2014). Furthermore, RyR distribution is dynamic (Asghari et al., 2020) and spacing between CRUs becomes shortened in failing hearts (Chen-Izu et al., 2007). Future studies on the cellular and molecular mechanisms regulating CRU distribution dynamics within cardiac cells will clarify how CRU order/disorder contributes to cell physiological and pathological function. While our numerical simulations show clearly a notable effect of disorder in CRU positions on Ca release synchronization and spontaneous AP firing, the theoretical mechanisms of this synchronization merit further studies. Based on our numerical study one can envision that synchronization is happening as a critical phenomenon, with the criticality depending on model parameters, including spatial randomness of the CRUs. Possible approaches to study such systems with criticalities include a percolation phase transition or Ising formalism, similar to what has been recently suggested for RyR interactions via CICR within CRUs (Maltsev et al., 2017a; Maltsev et al., 2019).

### Conclusions

The rate of Ca release propagation is an important feature of both normal and abnormal Ca release signals in cardiac cells. Using numerical modeling here we show that disorder in CRU locations increases the synchronization of Ca release in SA node pacemaker cells. This impacts on their pacemaker function via NCX current accelerating diastolic depolarization. While the disorder increases the rate and robustness of spontaneous AP firing, it simultaneously decreases βAR stimulation effect and the low range of lower rates. Thus, spatial imperfection encodes functional perfection: the disorder determines the rate of Ca release propagation and thereby allows pacemaker cells to regulate their balance of robustness and flexibility. On the other hand, excessive disorder in CRU position is expected to limit the heart rate range that may contribute to the heart rate range decline with age and in disease.

## Supporting information

Movies for various types of disorder

## Acknowledgments

This research was supported in part by the Intramural Research Program of the National Institutes of Health, National Institute on Aging. A.V.M. acknowledges the support of the Royal Society University Research Fellowship UF160569.

## APPENDIX

## Detailed methods

## 1. General description of the model

In the present study we performed a major update of our previous Ca release unit (CRU)-based numerical model of a central rabbit SAN cell (Maltsev et al., 2011; Maltsev et al., 2013). The model formulations for cell membrane currents are adopted from 2009 Maltsev-Lakatta model (Maltsev and Lakatta, 2009) that, in turn, stems from 2002 Kurata et al. (Kurata et al., 2002). The local Ca release (LCR) in the model is approximated at the scale of an individual CRU that represents a cluster of Ca release channels (ryanodine receptors, RyR) embedded in the junctional sarcoplasmic reticulum (JSR), i.e. a Ca store located in close proximity to cell surface membrane, with only 20 nm of separation via a dyadic space. Each JSR is diffusely linked to the network of free sarcoplasmic reticulum (FSR) that uptakes Ca from cytosol via SERCA pumping. Individual release channels are not modelled here, but we translate recent findings of RyR studies to the CRU level to introduce respective spark activation and termination mechanisms. All CRUs are identical and located in an equally-spaced square grid under the cell membrane. The model allows each CRU position to vary around its original square lattice position to generate a variety of intermediate distributions with various degrees of disorder from perfect square lattice to uniformly random. The specific aims of the present study were to update the model and to investigate how the disorder in CRU positions influences SAN cell function.

## 2. Cell geometry, compartments, voxels and membrane patches

We model a small SAN cell shaped as a cylinder of 53.28 μm in length and 6.876 μm in diameter. The cell membrane electrical capacitance of 19.8 pF is similar to that of 20 pF in Zhang et al. model (Zhang et al., 2000) of a central SAN cell. Details of local Ca dynamics under the cell membrane are simulated on a high-resolution (120 nm) square grid that divides the cell membrane and the submembrane space (dubbed subspace) into respective membrane patches and subspace voxels. Locations within the grid are defined by coordinates in the respective plane of the cell surface cylinder: along the cylinder (*x* axis) and around the cell cross-section (*y* axis). Our cell partitioning and respective voxel sizes to reproduce LCRs observed experimentally are schematically illustrated in the main text Fig. 1. To avoid special considerations at the cell borders, the ends of the cylinder are connected (yielding a torus). Our cell compartments and voxel structure are essentially similar to that in Stern et al. model (Stern et al., 2014) that approximates Ca dynamics in 3 dimensions at the single RyR scale. However, we limited the cell partition to only three nested layers of voxels of substantially different scales reflecting respective essential Ca cycling components and processes happening at these scales (described below).

## 2.1. Submembrane voxels

The first, most detailed level of approximation of local Ca dynamics is via small voxels of 120×120×20 nm under the entire cell membrane including the dyadic space (or cleft space) separating CRUs and the cell membrane. This thin layer of voxels describes local Ca release from individual CRUs, CRU-to-CRU interactions via Ca diffusion and CRU interactions with cell membrane (including *I*_*CaL*_, *I*_*NCX*_, *I*_*CaT*_, and *I*_*bCa*_). Ca currents were computed for each membrane patch (120×120 nm) to generate local Ca fluxes contributing to local Ca dynamics.

## 2.2. JSR level voxels (dubbed “ring” voxels)

The next approximation level of Ca dynamics is a deeper layer of voxels that have a larger size of 360×360×800 nm (*ΔxΔyΔr*) that includes the scale of JSR depth (60 nm). Each ring voxel has its cytoplasmic part and FSR part and some ring voxels are diffusively connected to JSRs (main text Fig. 1). While it appears that the geometric scale of ring voxel is of an order of magnitude larger than the JSR size, the actual FSR volume within each ring voxel is comparable with the JSR volume (described below in details). Thus, this level of voxels describes the local dynamics of Ca transfer from FSR to JSR as well as enhanced local Ca pumping and diffusion fluxes due to close proximity to Ca release and Ca influx in its neighboring submembrane voxels.

## 2.3. The core

In contrast to Stern model (Stern et al., 2014), the rest of the cell in our model does not have further geometric partitioning and it is lumped to one compartment “the core”, where local Ca dynamics is less important. The core also has its cytosolic and FSR parts (as in ring voxels). Thus, it describes the bulk Ca uptake from cytosol to FSR and further accumulation, diffusion, and redistribution of the pumped and released Ca within the cell interior.

## 2.4. JSR

We place JSRs inside the respective ring voxels just below their outer side facing the cell membrane (main text Fig. 1). JSR volume (~7.8 attoliter) is comparable with the volume of a ring voxel (~91 attoliter) and the same volume cannot be occupied twice by different cell compartments. Therefore, volumes of the ring voxels are kept the same by their extending into the core by the exact volume that the JSR occupies at their outer side. Thus, the actual core volume was calculated as the volume of the cylinder core decreased by the volume of all JSR volumes. We simulated different degree of disorder of JSR positions by a random number generator within a Gauss distribution along *x* and *y* (centered at the square lattice vertices) with a given SD that was the same for *x* and *y* directions. The model scenario with uniformly random positioning of CRU centers was also generated by a random number generator but with equal probability to occur in any submembrane voxel. JSR overlaps are excluded, i.e. any two JSRs cannot occupy the same cell volume.

## 3. Ca CYCLING

## 3.1. Free SR (FSR)

As mentioned above, each ring voxel and the core is further partitioned into cytosol and FSR fractions. We modelled FSR as homogeneously distributed network within the cytosol. The cytosol fraction was set to 0.46 and FSR fraction to 0.035 (Stern et al., 2014). The remainder presumably contains myofilaments, mitochondria, nucleus, and other organelles. In turn, each FSR portion has capability to pump Ca locally from the respective cytosol portion of the same voxel, simulating local SERCA function. Because the submembrane voxels are extremely thin, only 20 nm depth, their contribution to Ca pumping and intra-SR diffusion are negligible and not modelled.

## 3.2. SR Ca pump

The SERCA pump is present uniformly throughout the cell, transferring Ca from the cytosolic to the FSR compartment of each voxel (of ring and core) with Ca uptake rate given by the reversible Ca pump formulation adopted from Shannon et al. (Shannon et al., 2004)

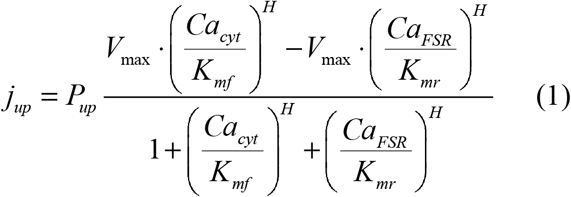

where *P*_*up*_ = 0.014 mM/ms, *K*_*mf*_ = 0.000246 mM, *K*_*mr*_ = 1.7 mM, and H = 1.787.

## 3.3. Ca diffusion within and among cell compartments

Ca diffusion fluxes between voxels within and among cell compartments are approximated by the first Fick’s law:

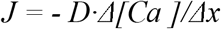

where *D* is a diffusion coefficient, and *Δ[Ca]/Δx* is Ca concentration gradients, i.e. *Δ[Ca]* is the concentration difference and *Δx* is the distance between the voxel centers.

The respective rate of change of [Ca] is defined as

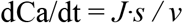

where *s* is the diffusion area sharing by voxels and *v* is the receiving volume. Thus

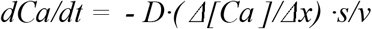

For any two diffusively interacting voxels with volumes *v*_*1*_ and *v*_*2*_, Ca dynamics is described by a set of differential equations:

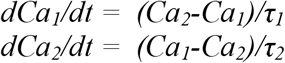

where *Ca*_*1*_ and *Ca*_*2*_ are Ca concentrations in the respective voxels and

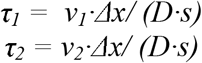

*τ*_*1*_ (or *τ*_*2*_, symmetrically) is the respective time constant of *Ca*_*1*_ change in time in a special case if the other compartment with volume *v*_*2*_ is substantially larger than *v*_*1*_ (i.e. *v*_*2*_ >> *v*_*1*_) and therefore *Ca*_*2*_ remains approximately constant. In general case, the set is analytically solved to the respective exponential decays:

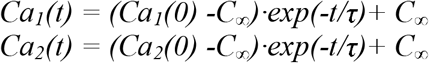

where *C*_∞_= *(Ca*_*1*_*(0)* ·*v1* + *Ca*_*2*_*(0)* · *v*_*2*_)/( *v*_*1*_ + *v*_*2*_) is equilibrium concentration (t = ∞) in both voxels defined by the matter conservation principle and *τ* =*τ*_*1*_·*τ*_*2*_/(*τ*_*1*_ + *τ*_*2*_) is the common time constant of the exponential decay of the system to reach the equilibrium. The respective Ca change from its initial value in voxel *v*_*1*_ over time is given as follows

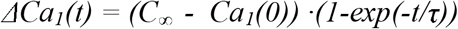

Then, by substituting C_∞_ we get:

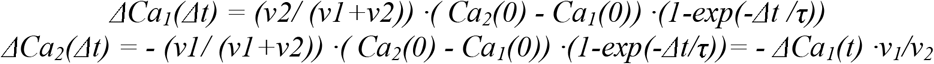

These formulations were used in all our computations of [Ca] changes for all neighboring voxels within and among cell compartments for the model integration for each time update *Δt* (during time tick or several time ticks for slower processes). In our computer algorithm we calculated the fractional Ca change (*FCC*) before the model run and used it simply as a scaling factor to determine actual [Ca] change from the difference in [Ca] between any two diffusely interacting voxels at the beginning of each integration step (from *t=0* to *t=Δt*). Thus,

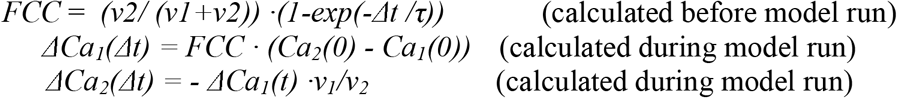

In the case of identical voxels, i.e. within subspace and within ring (i.e. when *v*_*1*_ = *v*_*2*_ and *τ*_*1*_= *τ*_*2*_) the formulations are simplified to:

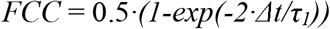

Note 1:

If *Δ*t/*τ*_*1*_ << 1 then *FCC* can be approximated (e.g. via respective Taylor series) as

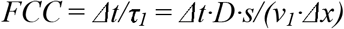

Further, if *v*_*1*_ can be described as *v_1_ = s · Δx,* e.g. for diffusion along the cell length (axis *x* in our model) *FCC* can be further simplified to

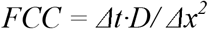

However, because *FCC* is calculated only once before the model run and does not carry any additional computational burden during actual simulations of Ca dynamics, we always used here the full approximation for the diffusion, i.e. more precise exponential decay, rather than a linear change over *Δt*. An advantage of this approach is that the model features more stable behavior in case we want to vary cell geometry, cell compartments, voxel sizes, or integration time (*Δt*).

Note 2:

We have only a fraction of cell volume occupied with cytosol (or FSR). However, the same fraction will be for *v* and *s* in *τ* formulations and it cancels. The volume ratios *v*_*2*_/(*v*_*1*_+*v*_*2*_) and *v*_*1*_/*v*_*2*_ remain also unchanged because the fraction factor also cancels. Thus, all above formulations with formal geometric volumes are also valid for fractional volumes, assuming that the fraction of cytosol (or FSR) is evenly distributed within the volume.

## 3.5. Junctional SR (JSR) and Ca diffusion between JSR and FSR

The JSRs are located in close proximity to the cell membrane separated only by the layer of submembrane voxels of 20 nm depth, representing the dyadic space. Thus, each CRU releases Ca (described below) into its neighboring subspace voxels (occupying the dyadic space). Each CRU is refilled with Ca locally from FSR network via a fixed diffusional resistance. In previous common pool models (Kurata et al., 2002; Maltsev and Lakatta, 2009) diffusion between JSR and FSR was described by a simple exponential transfer process with a fixed time constant (*τ*_*tr*_ =40 ms). Here we want to have a similar Ca transfer rate for each JSR, but locally. The *τ*_*tr*_ value in common pool models is for the whole cell FSR volume that is substantially larger than JSR volume. Here, in the local model, each JSR is linked diffusively to FSR. Depending on its position, JSR can be connected to one ring voxel, 2 voxels, or 4 voxels. To get the respective share of the Ca flux, we split and distribute the Ca diffusion flow into 9 elementary surface areas (120 nm x 120 nm) of the JSR (360×360 nm), each of which is connected to the respective ring voxel and transfer its Ca share with *τ* = 9·*40 ms* · *v*_*FSR*_/( *v*_*FSR*_ + *v*_*JSR*_) = 104.58 ms. Note: Because the FSR fraction of 0.035 within cell volume is rather small, each relatively large ring voxel of 360×360×800 nm (~91 attoliter) contains only a small FSR volume *v*_*FSR*_ = 3.186 attoliter. This is comparable (and even smaller) than the JSR volume in the model *v*_*JSR*_ = 7.776 attoliter.

## 3.6. Spark activation mechanism

Each CRU can be either in open or closed state. The capability of a given CRU to open, i.e. to generate a Ca spark, is controlled by its JSR Ca loading. Experimental and theoretical studies showed that sparks cannot be generated with SR Ca loading less than a certain critical level of about 300 μm (Zima et al., 2010; Veron et al., 2021). This critical level *Ca*_*jSR*_ is implemented in our mechanism of spark activation by prohibiting CRU firing while SR Ca loading remains below 300 μM (*CaSRfire*). When JSR is refilled with Ca above the *CaSRfire* level, it can open. The switch from close state to open state is probabilistic. The probability density for a given closed CRU to open is described by a power function of Ca concentration (*Ca*) in the dyadic space. The probability for the CRU to open during a short time interval *TimeTick* is given by

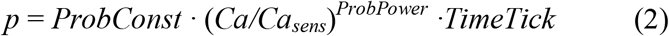

where *Ca*_*sens*_ = 0.15 μM sensitivity of CRU to Ca, *ProbConst =*0.00027 ms^-1^ is open probability rate at *Ca* = *Ca*_*sens*_, and *ProbPower* = 3 defines the cooperativity of CRU activation by cytosolic Ca. Each time tick our computer algorithm tries to activate a closed CRU by generating a random number within (0,1). If this number less than *p*, then the CRU opens. The Ca current amplitude, *I*_*spark*_, is defined by spark activation kinetics *a(t)*, RyR unitary current (*I*_*RyR,1mM*_, the current via a single RyR at 1 mM of Ca gradient), the number of RyRs residing in the JSR (*N*_*RyR*_), and concentration difference between inside and outside JSR:

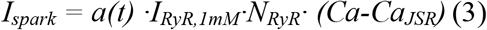

*N*_*RyR*_ is defined based on the surface area of JSR assuming a crystal-like structure for RyR positions separated by 30 nm. Thus, for our JSR *xy* area of 360 x 360 nm, we obtain 12 x 12 RyRs, i.e. *N*_*RyR*_ =144. *I*_*RyR,1mM*_ is to set 0.35 pA (Stern et al., 2014). Spark activation is described as a single exponential time-dependent process to tune spark rise time to about 5 ms (Fig. A1) close to that reported in the literature (from 4 to 8 ms).

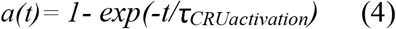

It is important to note that our new model spark initiation does not reflect an intrinsic time-dependent refractory process; rather we implement here the idea that spark can occur only when SR gets refilled to a critical level that the current amplitude of individual RyR can initiate regenerative CICR among neighboring RyRs (Zima et al., 2010; Stern et al., 2013; Veron et al., 2021). Thus, the implementation of the new spark activation mechanism controlled by SR Ca refilling represents a major advance of our model, because the spark activation timing is now predicted by the model. Of note, our previous CRU-based SAN cell models (Maltsev et al., 2011; Maltsev et al., 2013) implemented SR Ca refiling contribution phenomenologically via a fixed parameter, the restitution time that was directly taken from experimental measurements.

## 3.7. Spark termination mechanism

We also introduced a new spark termination mechanism that is based on the current knowledge in this research area (Laver et al., 2013; Stern et al., 2013; Maltsev et al., 2017a), i.e. a Ca spark is generated via CICR among individual RyRs within a CRU and it sharply terminates due to induction decay (Laver et al., 2013) or a phase transition (similar to that known in Ising model)(Maltsev et al., 2017a) when RyR current *I*_*RyR*_*(t)* becomes too small (due to JSR Ca depletion) to further support the CICR. The specific value of the critical current *I*_*spark_termination*_ is defined by the RyR interactions and beyond the capability of our CRU-based model. But the time when CICR wanes to the critical point is reflected by the amplitude of *I*_*spark*_*(t)* being comparable with *I*_*RyR*_*(t)* at a given JSR load, so that only one or a very few RyRs remain open at the termination time point when spark decays. Based on this logic, we tested a wide range of *I*_*spark_termination*_ to generate sparks of various durations and found that spark termination is well described with *I*_*spark_termination*_ set to a 0.175 pA. In our previous numerical model simulations and Ising theory of spark (Maltsev et al., 2017a) spark termination happens when SR level depletes to a critical level of about 0.1 to 0.15 mM (for RyR clusters from 9×9 to13×13). Unitary RyR current *I*_*RyR*_ becomes very small at these SR levels, namely within the range of 0.035 pA to 0.0525 pA, respectively, assuming *I*_*RyR*_ at 1mM of SR Ca to be *I*_*RyR,1mM*_ = 0.35 pA (as in (Stern et al., 2013)). Thus, our chosen critical spark amplitude of 0.175 pA is reasonable for spark termination time point, reflecting only 2 or 3 remaining open RyR channels (of 144 total in our SAN cell model). As the SR continues to be further depleted of Ca, these few open channels cannot support any longer CICR within the CRU, and spark undergoes a sharp termination phase transition that is also in line with our numerical simulations of spark dynamics (Figures 1 and 3 in (Maltsev et al., 2017a)). It is important to note that the spark termination mechanism does not include any intrinsic time-dependent inactivation, but simply reflects the drop of local Ca gradient over the JSR to the critical point that cannot sustain CICR among RyRs within the CRU (Laver et al., 2013; Stern et al., 2013; Maltsev et al., 2017a)).

## 3.8. Ca buffering

Cytosolic Ca is buffered by calmodulin (0.045 mM) throughout the cell: submembrane voxels, ring voxels, and the core. Each JSR features Ca buffering with calsequestrin (30 mM).

## 3.9. Summary of equations of local Ca dynamics

## In a submembrane voxel

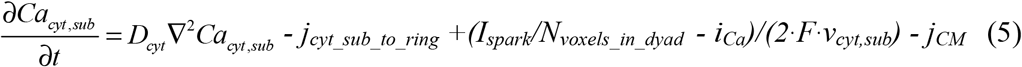

*I*_*spark*_ in voxels outside dyadic space is absent. Each CRU releases Ca (given by *I*_*spark*_) into its dyadic space. *I*_*spark*_ is evenly distributed among submembrane voxels of the dyadic space (*N*_*voxels_in_dyad*_ = *9*). *I*_*spark*_ is given in Equation 3 and it is positive, i.e. increasing [Ca] in the submembrane voxel. *i*_*Ca*_ is the sum of local Ca transmembrane currents (described in details below in Electrophysiology section) via the membrane patch facing this submembrane voxel:

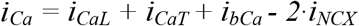

*I*_*CaL*_ is included in this equation only for submembrane voxels facing a CRU (*I*_*CaL*_ is injected into the 9 subspace voxels of the respective dyadic space). The local Ca currents *i*_*CaL*_, *i*_*CaT*_, *i*_*bCa*_ have inward direction and (by convention) are defined as negative. Therefore, the minus sign before *i*_*Ca*_ in Equation 5 ensures positive change in [Ca] in the submembrane voxel by respective Ca influx. During diastole local *i*_*NCX*_ also flows inwardly, but NCX exchanges 1 Ca ion to 3 Na ions. Hence, the inward *i*_*NCX*_ generates a Ca efflux. That is why it has a different sign.

## In a voxel of ring layer

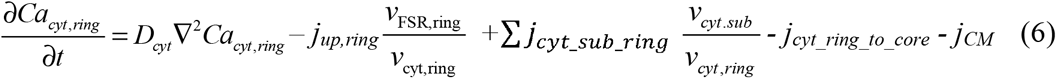

∑*j*_*cyt_sub_ring*_ is the sum of diffusion fluxes from neighboring smaller submembrane voxels

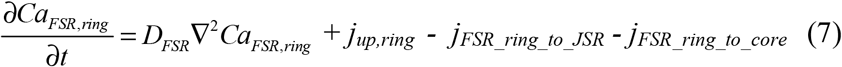

The SERCA uptake flux *j*_*up*_ is given by Equation 1.

## In a given JSR

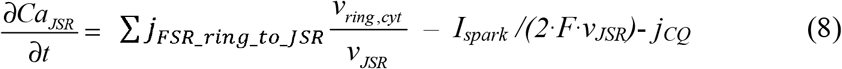

where *I*_*spark*_ is given by Equation 3.

∑*j*_*FSR_ring_to_JSR*_ is the sum of diffusion fluxes between JSR and respective FSR parts of the neighboring ring voxels. *j*_*CQ*_ is Ca flux of Ca buffering by calsequestrin:

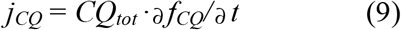

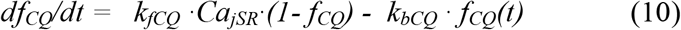

## In the core

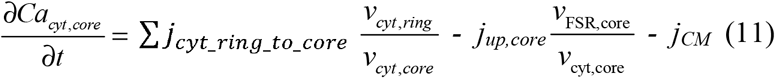

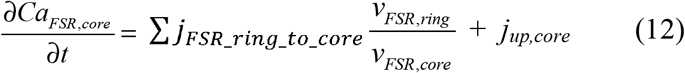

where ∑*j*_*cyt_ring_to_core*_ and ∑*j*_*FSR_ring_to_core*_ are the respective sums of diffusion fluxes in cytosol and FSR of all ring voxels.

## In any cytoplasmic voxel (subspace, ring, and the core)

*j*_*CM*_ is Ca flux of Ca buffering by calmodulin:

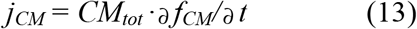

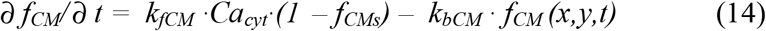

## 4. ELECTROPHYSIOLOGY

Electrophysiological formulations for cell membrane currents are adopted from (Maltsev and Lakatta, 2009). Major changes of the model include introduction of local Ca currents and modulation of local currents by local Ca. To introduce local currents and local modulation by Ca, the cell membrane is partitioned into small patches, with each patch facing its respective subspace voxel. We also omitted sustained inward current *I*_*st*_ and background Na current *I*_*bNa*_ because thus far the molecular identities for these currents have not been found and these currents are likely produced by NCX or other currents (Lakatta et al., 2010). We adopted *I*_*NCX*_ density and Ca-dependent *I*_*CaL*_ inactivation for more realistic modulation by much higher local Ca concentrations under the cell membrane predicted by our local Ca control models in the present study (Fig. A1) and previous models (Maltsev et al., 2011; Maltsev et al., 2013; Stern et al., 2014; Maltsev et al., 2017b) reaching >10 μM vs. common pool models (Kurata et al., 2002; Maltsev and Lakatta, 2009) predicting [Ca] in subspace only in sub-μM range in a bulk “subspace” compartment during diastole and 1-2 μM during Ca transient peak.

## 4.1 Fixed ion concentrations, mM

*Ca*_*o*_ = *2*: Extracellular Ca concentration.

*K*_*o*_ = *5.4*: Extracellular K concentration.

*K*_*i*_ =*140*: Intracellular K concentration.

*Na*_*o*_ = *140*: Extracellular Na concentration.

*Na*_*i*_ =*10*: Intracellular Na concentration.

## 4.2. The Nernst equation and electric potentials, mV

*E*_*X*_ = *(RT/F)* · *ln([X]*_*o*_/*[X]*_*i*_) = *E*_*T*_ · *ln([X]*_*o*_/*[X]*_*i*_), where

*F* = *96485 C/M* is Faraday constant,

*T* = *310.15 K°* is absolute temperature for 37°C,

*R* = *8.3144 J/(M·K°)* is the universal gas constant,

*E*_*T*_ is “*RT/F*” factor = 26.72655 mV, and [X]_o_ and [X]_i_ are concentrations of an ion “X” out and inside cell, respectively

*E*_Na_ = *E*_T_ · ln(Na_o_/Na_i_): Equilibrium potential for Na

*E*_*K*_ = *E*_*T*_ · *ln(K*_*o*_/*K*_*i*_): Equilibrium potential for K

*E*_*Ks*_ = *E*_*T*_ · *ln[(K*_*o*_ + *0.12*· *Na*_*o*_)/(*K*_*i*_ + *0.12* · *Na*_*i*_)]: Reversal potential of *I*_Ks_

*E*_*CaL*_ = *45*: Apparent reversal potential of *I*_*CaL*_

*E*_*CaT*_ = *45*: Apparent reversal potential of *I*_*CaT*_

## 4.3. Membrane potential, *V*_*m*_

Net membrane current determines time derivative of the membrane potential.

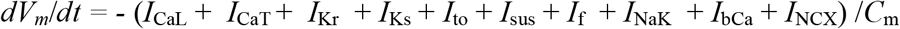

## 4.4. Formulation of cell membrane ion currents

Kinetics of ion currents are described by gating variables (described below) in respective differential equations

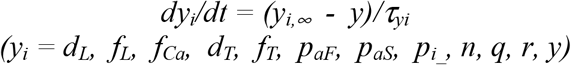

*τ*_*yi*_: time constant for a gating variable *y*_*i*_.

*α*_*yi*_ and *β*_*yi*_: opening and closing rates for channel gating.

*y*_*i*_,_∞_: steady-state curve for a gating variable *y*_*i*_.

## L-type Ca current *(I*_*CaL*_)

The whole cell *I*_*CaL*_ is calculated as a sum of local currents *i*_*CaL,i*_ in each dyadic space.

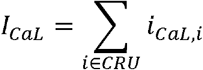

This reflects reports that L-type Ca channels are colocalized with RyRs (Christel et al., 2012). Thus, the whole cell maximum *I*_*CaL*_ conductance (*g*_*CaL*_) is distributed locally and equally among dyadic spaces:

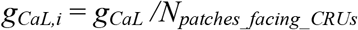

The respective local *i*_*CaL,i*_ is calculated based on formulations of Kurata et al. (Kurata et al., 2002) and subsequent modifications in our common pool models (Maltsev and Lakatta, 2009, 2010, 2013), but now its Ca-dependent inactivation is determined by local subspace [Ca] (*Ca*_*sub,i*_):

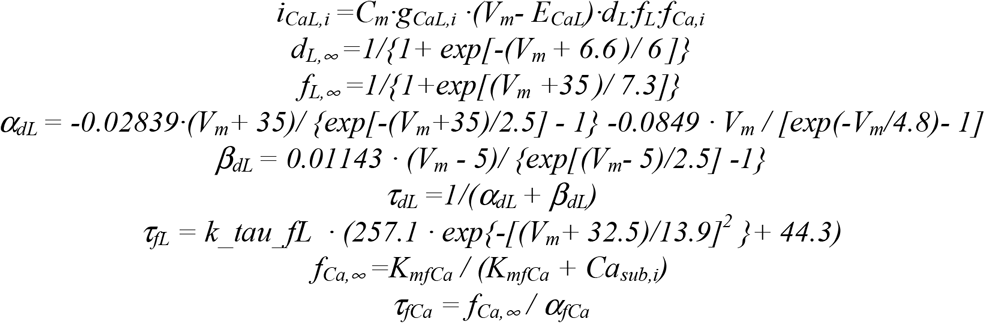

Ca-dependent *I*_CaL_ inactivation is described by parameters *K*_mfCa_ and *α*_fCa_. With *K*_mfCa_ of 0.35 μM in previous common pool models, our local Ca model of SAN cell did not work (not enough *I*_*CaL*_ was available to generate normal APs), because *I*_CaL_ inactivation was too sensitive to Ca and a major part of *I*_*CaL*_ was inactivated in diastole by diastolic LCRs whose amplitudes reach tens and hundreds of μM. Thus, we adopt Ca-dependent inactivation of *I*_*CaL*_ for more realistic modulation by much higher local Ca concentrations by setting *K*_mfCa_ to 30 μM. We also set the midpoint of *I*_*CaL*_ activation to −6.6 mV as in SAN cell models of Wilders et al. (Wilders et al., 1991) and Dokos et al. (Dokos et al., 1996).

## T-type Ca current *(I*_*CaT*_)

It is based on formulations of Demir et al.(Demir et al., 1994) and modified by Kurata et al.(Kurata et al., 2002).

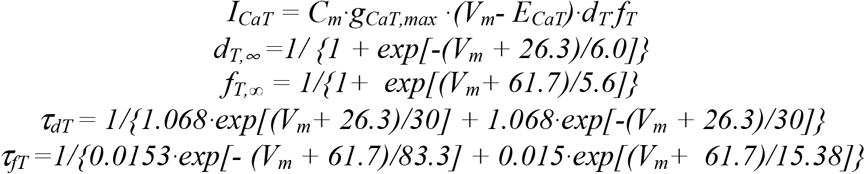

The whole cell *I*_*CaT*_ was evenly distributed over cell membrane patches to generate respective homogeneous Ca influx. In each submembrane *i-th* voxel *I*_*CaT,i*_ = *I*_*CaT*_*/N*_*voxels*_

## Rapidly activating delayed rectifier K^+^ current (*I*_Kr_)

It is based on formulations of Zhang et al. (Zhang et al., 2000), further modified by Kurata et al. (Kurata et al., 2002).

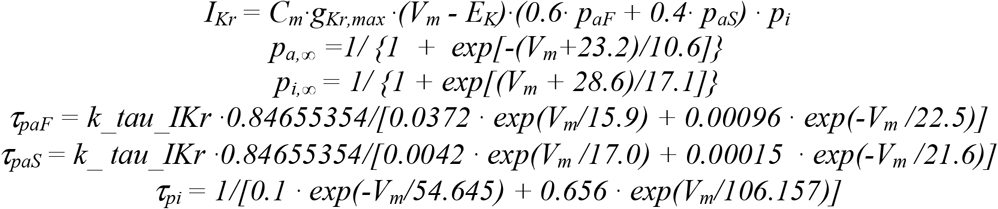

## Slowly activating delayed rectifier K^+^ current (*I*_Ks_)

It is based on formulations of Zhang et al. (Zhang et al., 2000).

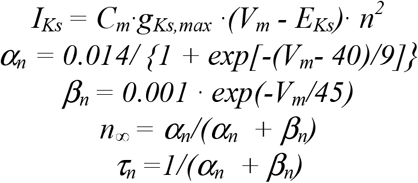

## 4-aminopyridine-sensitive currents (*I*_4AP_ =*I*_to_ + *I*_sus_)

It is based on formulations of Zhang et al. (Zhang et al., 2000).

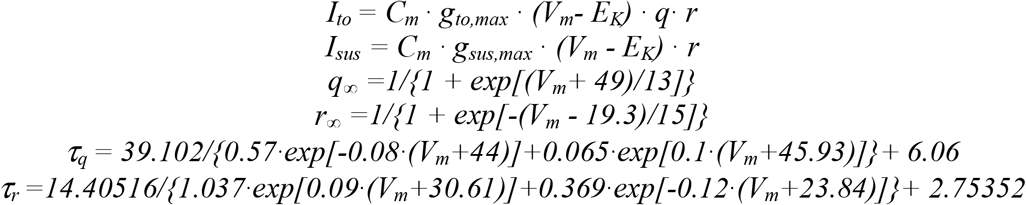

## Hyperpolarization-activated, “funny” current (*I*_*f*_)

It is based on formulations of Wilders et al.(Wilders et al., 1991) and Kurata et al.(Kurata et al., 2002).

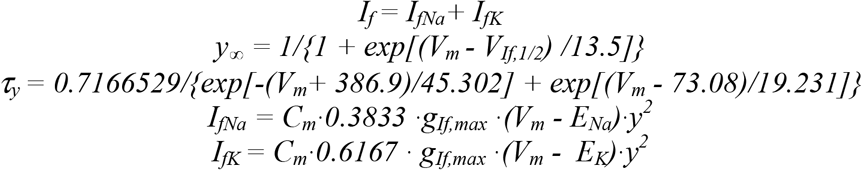

## Na^+^-K^+^ pump current (*I*_*NaK*_)

It is based on formulations of Kurata et al.(Kurata et al., 2002), which were in turn based on the experimental work of Sakai et al.(Sakai et al., 1996) for rabbit SAN cell.

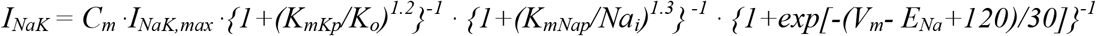

## Ca- background current (*I*_*bCa*_)

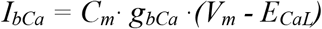

The whole cell *I*_*bCa*_ was evenly distributed over cell membrane to generate respective homogeneous Ca influx. In each submembrane *i-th* voxel *I*_*bCa,i*_ = *I*_*bCa*_*/N*_*voxels*_

## Na-Ca exchanger current (*I*_NCX_)

It is based on original formulations from Dokos et al. (Dokos et al., 1996). *I*_*NCX*_ is modulated by local Ca and therefore the whole cell *I*_*NCX*_ was calculated as a sum of local currents *I*_*NCX,i*_ in respective membrane patches facing each submembrane voxel. Thus,

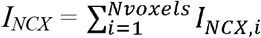

For membrane voltage *V*_*m*_ in each *i*-th membrane patch with subspace *Ca*_*sub,i*_, the respective local NCX current (*I*_*NCX,i*_) was calculated as follows:

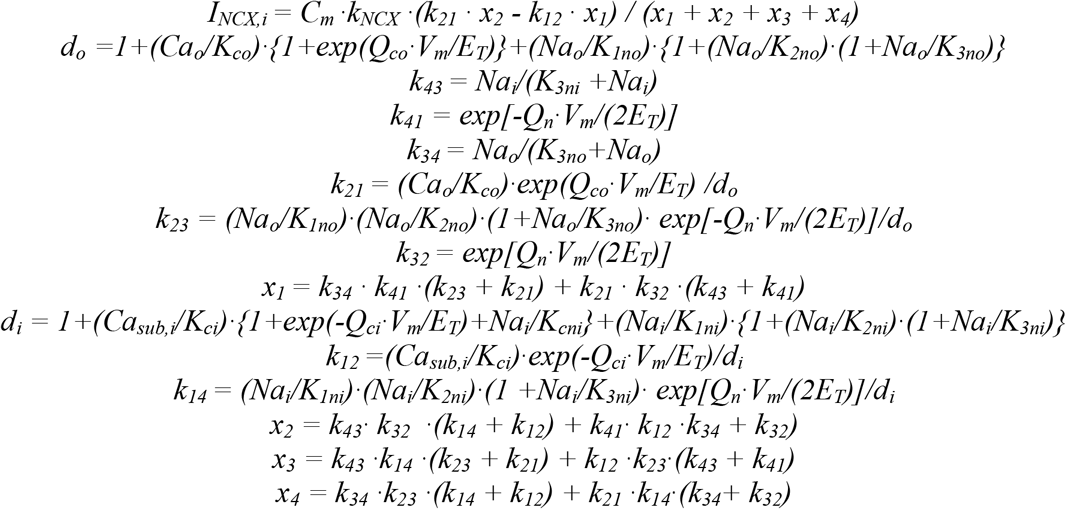

## 5. Initial values

Initial values for electrophysiology were as follows:

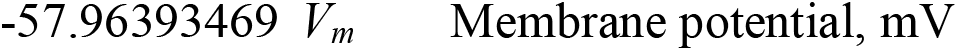

|  |  |  |
| --- | --- | --- |
| 0.000584546 | $d_L$ | $I_{CaL}$ voltage-dependent activation |
| 0.86238125 | $f_L$ | $I_{CaL}$ voltage-dependent inactivation |
| 0.71139592 | $f_{Ca}$ | $I_{CaL}$ Ca-dependent inactivation (local) |
| 0.144755091 | $p_{aF}$ | $I_{Kr}$ fast activation |
| 0.453100577 | $p_{aS}$ | $I_{Kr}$ slow activation |
| 0.849409822 | $p_{i-}$ | $I_{Kr}$ inactivation |
| 0.026460041 | $n$ | $I_{Ks}$ activation |
| 0.113643187 | $y$ | $I_f$ activation |
| 0.005043934 | $d_T$ | $I_{CaT}$ activation |
| 0.420757825 | $f_T$ | $I_{CaT}$ inactivation |
| 0.694241314 | $q$ | $I_{to}$ inactivation |
| 0.005581317 | $r$ | $I_{to}$ and $I_{sus}$ activation |

Initial values for Ca dynamics were as follows:

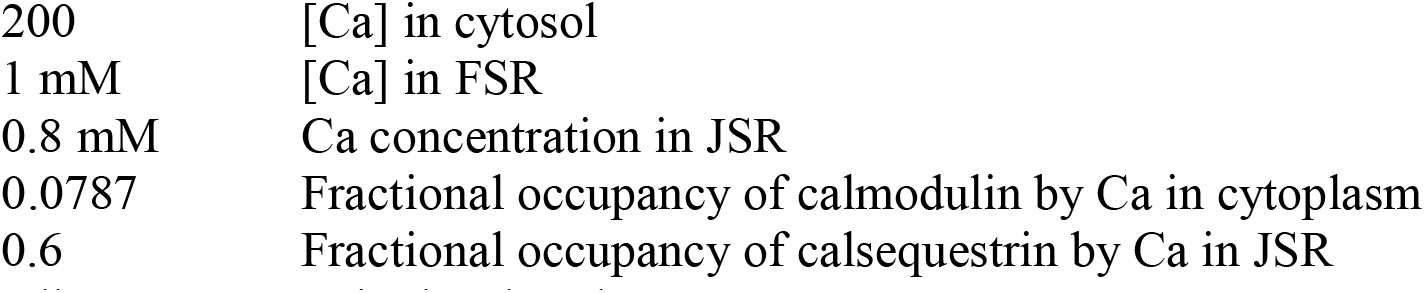

|  |  |
| --- | --- |
| 200 | [Ca] in cytosol |
| 1 mM | [Ca] in FSR |
| 0.8 mM | Ca concentration in JSR |
| 0.0787 | Fractional occupancy of calmodulin by Ca in cytoplasm |
| 0.6 | Fractional occupancy of calsequestrin by Ca in JSR |

All CRUs are set in the closed state

## 6. Summary of model parameters

## Fixed ion concentrations

*Ca*_*o*_=2 mM: Extracellular [Ca]

*K*_*o*_=5.4 mM: Extracellular [K]

*Na*_*o*_=140 mM: Extracellular [Na]

*K*_*i*_=140 mM: Intracellular [K]

*Na*_*i*_=10 mM: Intracellular [Na]

## Membrane currents

*E*_CaL_ = 45 mV: Apparent reversal potential of *I*_*CaL*_

*g*_*CaL*_ = 0.464 nS/pF: Conductance of *I*_*CaL*_

*K*_*mfCa*_ = 0.03 mM: Dissociation constant of Ca -dependent *I*_CaL_ inactivation

*beta*_fCa_ = 60 mM^-1^ · ms^-1^: Ca association rate constant for *I*_*CaL*_.

*alfa*_fCa_ = 0.021 ms^-1^: Ca dissociation rate constant for *I*_CaL_

*k_tau_fL* = 0.5: Scaling factor for *tau_fL* used to tune the model

*E*_CaT_ = 45: Apparent reversal potential of *I*_*CaT*_, mV

*g*_*CaT*_ = 0.1832 nS/pF: Conductance of *I*_*CaT*_

*g*_*If*_ = 0.105 nS/pF: Conductance of *I*_*f*_

*V*_*If,1/2*_ = −64: half activation voltage of *I*_f_, mV

*g*_*Kr*_ = 0.05679781 nS/pF: Conductance of delayed rectifier K current rapid component

*k_ tau_IKr* = 0.3: scaling factor for *tau_paF* and *tau_paS* used to tune the model

*g*_*Ks*_ = 0.0259 nS/pF: Conductance of delayed rectifier K current slow component

*g*_*to*_ = 0.252 nS/pF: Conductance of 4-aminopyridine sensitive transient K^+^ current

*g*_sus_ = 0.02 nS/pF: Conductance of 4-aminopyridine sensitive sustained K^+^ current

*I*_NaKmax_ = 1.44 pA/pF: Maximum Na^+^/K^+^ pump current

*K*_mKp_ = 1.4 mM: Half-maximal *K*_o_ for *I*_NaK_.

*K*_mNap_ = 14 mM: Half-maximal *Na*_i_ for *I*_NaK_.

*g*_bCa_ = 0.003 nS/pF: Conductance of background Ca current,

*k*_NCX_ = 48.75 pA/pF: Maximumal amplitude of *I*_*NCX*_

## Dissociation constants for NCX

*K*_1ni_ = 395.3 mM: intracellular Na binding to first site on NCX

*K*_2ni_ = 2.289 mM: intracellular Na binding to second site on NCX

*K*_3ni_ = 26.44 mM: intracellular Na binding to third site on NCX

*K*_1no_ = 1628 mM: extracellular Na binding to first site on NCX

*K*_2no_ = 561.4 mM: extracellular Na binding to second site on NCX

*K*_3no_ = 4.663 mM: extracellular Na binding to third site on NCX

*K*_ci_ = 0.0207 mM: intracellular Ca binding to NCX transporter

*K*_co_ = 3.663 mM: extracellular Ca binding to NCX transporter

*K*_cni_ = 26.44 mM: intracellular Na and Ca simultaneous binding to NCX

## NCX fractional charge movement

*Q*_ci_= 0.1369: intracellular Ca occlusion reaction of NCX

*Q*_co_=0: extracellular Ca occlusion reaction of NCX

*Q*_n_= 0.4315: Na occlusion reactions of NCX

## Ca buffering

*k*_*bCM*_ = 0.542 ms^-1^: Ca dissociation constant for calmodulin

*k*_*fCM*_ = 227.7 mM^-1^· ms^-1^: Ca association constant for calmodulin

*k*_*bCQ*_ = 0.445 ms^-1^: Ca dissociation constant for calsequestrin

*k*_*fCQ*_ = 0.534 mM^-1^· ms^-1^: Ca association constant for calsequestrin

*CQ*_*tot*_ = 30 mM: Total calsequestrin concentration

*CM*_*tot*_ = 0.045 mM: Total calmodulin concentration

## SR Ca ATPase function

*K*_*mf*_ = 0.000246 mM: the cytosolic side *K*_*d*_ of SR Ca pump

*K*_*mr*_ = 1.7 mM: the lumenal side *K*_*d*_ of SR Ca pump

H = 1.787: cooperativity of SR Ca pump

*P*_*up*_ = 0.014 mM/ms: Maximal rate of Ca uptake by SR Ca pump

## CRU (Ca release and JSR)

CRU_Casens = 0.00015 mM: sensitivity of Ca release to Casub

CRU_ProbConst = 0.00027 ms^-1^: CRU open probability rate at Casub=CRU_Casens

CRU_ProbPower = 3: Cooperativity of CRU activation by Casub

CaJSR_spark_activation = 0.3 mM: critical JSR Ca loading to generate a spark (CRU can open)

Ispark_Termination = 0.175 pA: critical minimum *I*_*spark*_ triggering spark termination (CRU closes) Ispark_activation_tau_ms = 80 ms: Time constant of spark activation (a)

Iryr_at_1mM_CaJSR = 0.35 pA: unitary RyR current at 1 mM delta Ca

RyR_to_RyR_distance_mkm = 0.03 μm: RyR crystal grid size in JSR

JSR_depth_mkm = 0.06 μm: JSR depth

jSR_Xsize_mkm = 0.36 μm: JSR size in x

jSR_Ysize_mkm = 0.36 μm: JSR size in y

jSR_to_jSR_X_mkm = 1.44 μm: JSR crystal grid size in x

jSR_to_jSR_Y_mkm = 1.44 μm: JSR crystal grid size in y

## Cell geometry, compartments, and voxels

*L*_cell_ = 53.28 μm: Cell length

*r*_cell_ = 3.437747 μm: Cell radius

*C*_*m*_ = 19.80142 pF: membrane electrical capacitance of our cell model with 0.0172059397937 pF/μm^2^ specific membrane capacitance calculated from 2002 Kurata et al. model (Kurata et al., 2002) for its 32 pF cylinder cell of 70 μm length and 4 μm radius.

0.12 μm: the grid size

0.12 μm: Submembrane voxel size in *x*

0.12 μm: Submembrane voxel size in *y*

0.02 μm: Submembrane voxel size in *r*

0.36 μm: Ring voxel size in *x*

0.36 μm: Ring voxel size in *y*

0.8 μm: Ring voxel size in *r*

0.46: Fractional volume of cytosol

0.035: Fractional volume of FSR

## Ca diffusion

*Dcyt* = 0.35 μm^2^/ms: Diffusion coefficient of free Ca in cytosol

*D*_*FSR*_ = 0.06 μm^2^/ms: Diffusion coefficient of free Ca in FSR

## Model Integration

TimeTick = 0.0075 ms

## 7. *g*_*CaL*_ sensitivity analysis

For square lattice and uniform random distributions of CRUs we performed sensitivity analysis for *g*_*CaL*_ from its basal value of 0.464 nS/pF (100%) down to 0.2552 nS/pF (55%) shown by magenta band in the main text Fig. 7 A with a step of 0.0232 pA/pF (5%). The original data of this analysis in the form of *V*_*m*_ time series are given Fig. A2 for each CRU distribution.

## 8. Simulations β AR stimulation effect

Effect of βAR stimulation was modelled essentially as we previously reported (Maltsev and Lakatta, 2010) by increasing *I*_*CaL*_, *I*_*Kr*_, *I*_*f*_, and Ca uptake rate by FSR via SERCA pumping. Specifically whole cell maximum *I*_*CaL*_ conductance *g*_*CaL*_ was increased by a factor of 1.75 from 0.464 to 0.812 nS/pF; whole cell maximum *I*_*Kr*_ conductance *g*_*Kr*_ was increased by a factor of 1.5 from 0.05679781 to 0.085196715 nS/pF; the midpoint of *I*_*f*_ activation curve was shifted to more depolarized potential by 7.8 mV from −64 mV to −56.2 mV; and the maximum Ca uptake rate *P*_*up*_ was increased by a factor of 2 from 14 mM/s to 28 mM/s. The original data in the form of intervalograms are given Fig. A3 for each CRU distribution in basal state and in during βAR stimulation

## Figure legends

**Figure A1.**
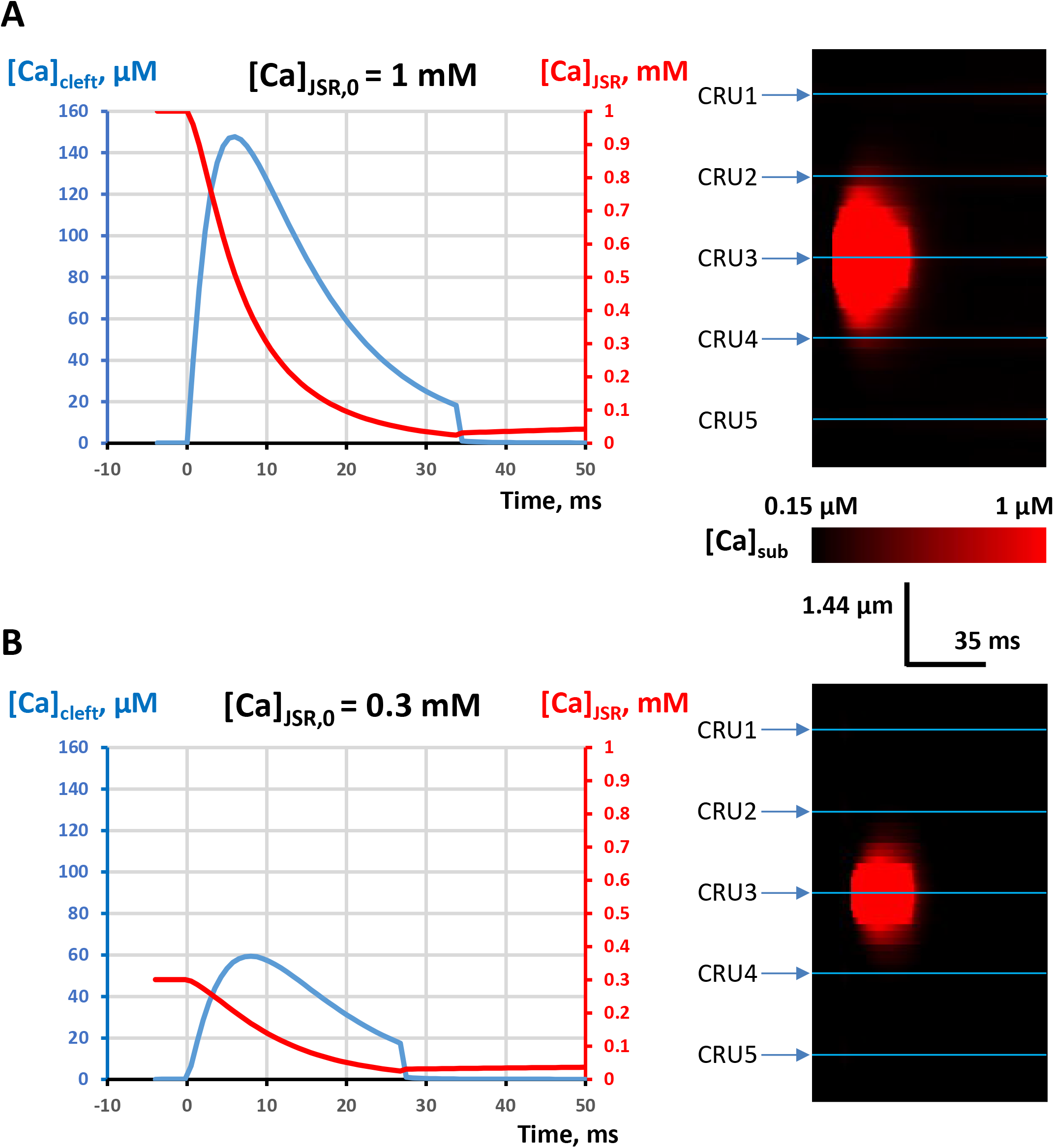
Representative Ca sparks generated by our CRU model at two different initial JSR Ca loading 1 mM (A) and 0.3 mM (B). Left panels: Overlapped time series of [Ca] in dyadic cleft (blue line) and [Ca] in JSR (red line). Right panels: Respective line-scan images of the sparks. The spark at 0.3 mM of JSR Ca loading had a lower amplitude and duration. To better illustrate Ca spread, the color map in line-scan images is saturated at [*Ca]*_*cleft*_ = 1 μM. CRU positions for square lattice configuration are shown by blue lines. The spark was generated by CRU3 and the images show that its outskirt could reach neighbouring CRUs (CRU2 and CRU4) at 1 mM JSR Ca loading, but not at 0.3 mM (i.e. at the *[Ca]*_*JSR*_ threshold of spark activation). The membrane potential was −60 mV.

**Figure A2.**
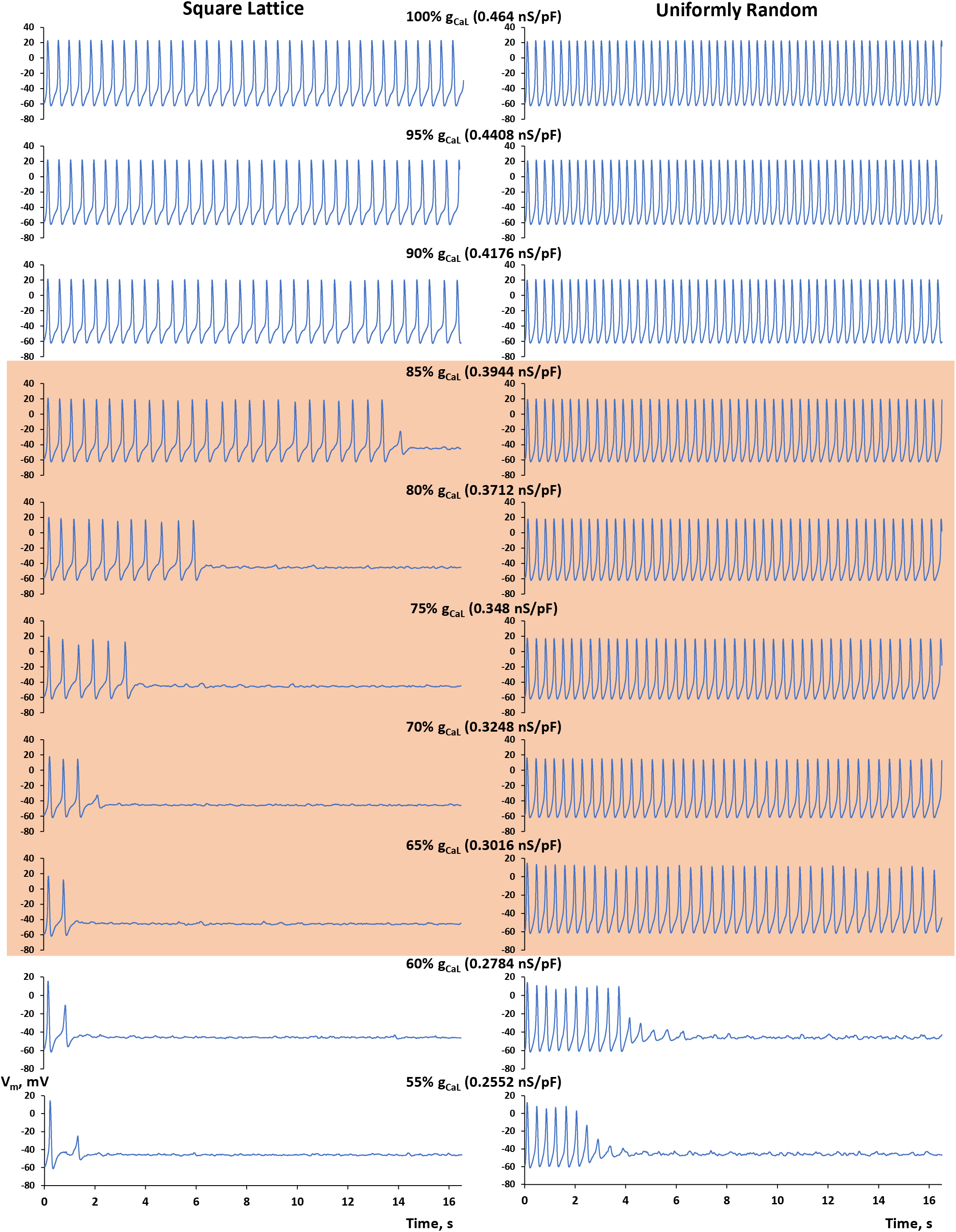
SAN cell with uniformly random distribution of CRUs features more robust spontaneous AP firing (right panels) vs. that with a square lattice distribution of CRUs (left panels). Shown are simulated AP traces in our *g*_*CaL*_ sensitivity analysis in which *g*_*CaL*_ gradually decreased from 100% to 55% of its basal state value of 0.464 nS/pF. Specific *g*_*CaL*_ values are shown at the top of each panel. Red shade shows the AP safety margin (functional reserve) that a SAN cell can utilize to increase its robust function via redistribution of its CRU locations to increase the presence of noise (i.e. natural CRU clustering).

**Figure A3.**
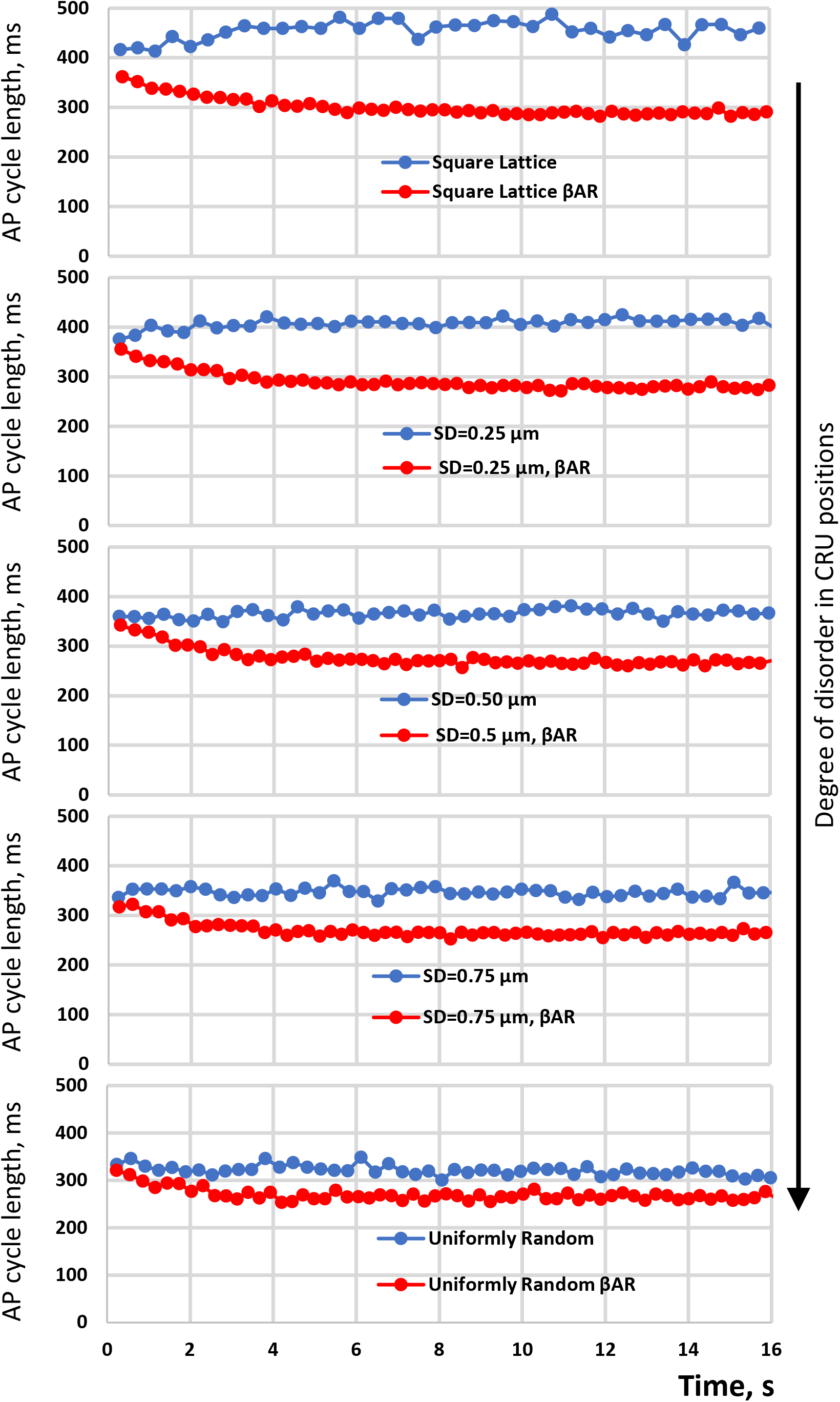
Effect of βAR stimulation wanes as disorder in CRU positions increases. Shown are intervalograms for spontaneous AP firing for 16 s of simulation time starting from the identical initial conditions in basal state (blue plots) and in the presence of βAR stimulation (red plots). Note: βAR stimulation shortens the cycle lengths towards about the same level within a narrow range of 291 ms to 266 ms, i.e. βAR stimulation unifies spontaneous AP firing towards a higher common rate.

## Movie legends

**Movie 1:**

**Square lattice distribution of CRUs**. Simulation of Ca dynamics in submembrane space (*[Ca]*_*sub*_) during 1s. Note mainly individual sparks in diastole. [*Ca]*_*sub*_ is coded from 0.15 μM (black) to 10 μM (saturation) by a color scheme shown at the bottom of Fig. 6; open CRUs are shown by white dots. Closed CRUs in refractory period are shown by blue dots; closed reactivated CRUs (available to fire) are shown by green dots. JSR Ca level is coded by respective shade (white, blue or green) with a saturation level set at 0.3 mM. Simulation time and membrane potential (*V*_*m*_) are shown in the top left corner.

**Movie 2:**

**Perturbed square lattice distribution of CRUs with SD=0.25**μ**m**. Sizes of LCRs increase during diastolic depolarization via propagating CICR. See Movie 1 legend for color description and other details.

**Movie 3:**

**Perturbed square lattice distribution of CRUs with SD=0.5**μ**m**. Sizes of LCRs further increase during diastolic depolarization via propagating CICR. See Movie 1 legend for color description and other details.

**Movie 4:**

**Perturbed square lattice distribution of CRUs with SD=0.75**μ**m**. Sizes of LCRs further increase during diastolic depolarization via propagating CICR. See Movie 1 legend for color description and other details.

**Movie 5:**

**Uniformly random distribution of CRUs**. Sizes of LCRs further increase during diastolic depolarization via propagating CICR. See Movie 1 legend for color description and other details.

